# Unique pore architecture underlies constitutive gating of human retinal TRPM1

**DOI:** 10.64898/2026.02.18.706614

**Authors:** Mansi Sharma, KV Nageswar, Vikesh Kumar, Ankur Chattopadhyay, Sristi Nanda, Nishtha Varshney, Rohit Chettri, Kirill A. Martemyanov, Appu Kumar Singh

## Abstract

Transient receptor potential melastatin 1 (TRPM1), a Ca^²⁺^-permeable nonselective cation channel essential for retinal ON bipolar cell signaling and night vision, and implicated in congenital night blindness, has remained structurally and functionally poorly characterized. Here we report the first cryo–electron microscopy structure of human TRPM1 in conducting state. Although the channel assembles as a tetramer, it adopts an unexpected clockwise domain-swapped pore module with rotational geometry inverse to that observed in previously characterized 6-TM tetrameric channels. This inverted topology is accompanied by extensive remodeling of the S5–P–S6 module, dilation of the selectivity filter, expansion of the central cavity, and splaying of S6 to form a wide intracellular gate. Single-channel recordings reveal constitutive activity consistent with the conductive state captured. Together, these findings uncover a new 6-TM fold in tetrameric channel and provide a framework for understanding TRPM1 gating and disease-associated dysfunction.

## INTRODUCTION

Members of the transient receptor potential (TRP) superfamily predominantly function as a tetrameric cation-conducting channels that play crucial roles in sensory physiology, including touch, pain, thermosensation, and vision. Transient receptor potential melastatin 1 (TRPM1) is a calcium-permeable, nonselective cation channel with essential function in visual signal transduction and pigmentation(*1*). In the retina, TRPM1 enables synaptic communication of photoreceptors. It is expressed in postsynaptic ON bipolar cells, mediating their depolarization in response to a light-evoked decrease in presynaptic glutamate released from photoreceptor synaptic terminals. Since ON bipolar neurons are the only synaptic partners of rod photoreceptors, TRPM1 is indispensable for rod-mediated scotopic vision under dim light conditions (*2*). Accordingly, knockout of TRPM1 in mice abolishes scotopic vision (REF). Mice lacking TRPM1 also have cognitive and behavioral abnormalities (*3*). In humans, loss-of-function mutations in TRPM1 cause congenital stationary night blindness (*4*). Additionally, decreased TRPM1 expression has been associated with melanoma progression and metastasis, highlighting the physiological and clinical significance of this channel (*5*). Despite being the founding member of the TRPM family, the structural basis of TRPM1 assembly and pharmacological modulation has remained largely unknown, greatly limiting mechanistic insight and therapeutic development into TRPM1 channelopathies. Therefore, a detailed understanding of TRPM1 structure is essential for defining its gating mechanisms and enabling structure-guided therapeutic strategies targeting TRPM1-associated visual and pigmentary disorders.

Tetrameric ion channels evolved from simple two–transmembrane pore modules into the canonical six–transmembrane (6-TM) architecture, in which an S1–S4 voltage-sensor or voltage sensor–like domain is functionally coupled to an S5–P–S6 pore module. In most structurally characterized 6-TM channels, including voltage-gated sodium, calcium, and potassium channels, the pore domain is arranged in a counterclockwise orientation relative to the voltage-sensing domain when viewed from the extracellular side, typically resulting in a domain-swapped architecture (*6–9*). More recently, non–domain-swapped configurations have been identified, often associated with a shortened S4–S5 linker that maintains the pore within the same subunit(*10–12*). TRP channels adopt 6-TM domain-swapped tetrameric structural framework but are distinguished by large cytoplasmic N- and C-terminal assemblies that integrate regulatory and signaling inputs(*13, 14*). Structures of TRPM2, TRPM3, TRPM4, TRPM7, and TRPM8 have established shared mechanisms of ion permeation and gating within the subfamily (*15, 16*). In contrast, TRPM1 exhibits distinct mode of physiological regulation: in retinal ON bipolar cells, it is tonically active in darkness and is suppressed by mGluR6 signaling through the G protein Gαo and βγ (*17, 18*). This specialized mode of regulation suggests that TRPM1 harbors structural features that diverge from established TRPM paradigms. Notably, although TRPM1 is most closely related to TRPM3, a previous structural study proposed an unexpected dimeric assembly for TRPM1, further highlighting uncertainties surrounding its quaternary organization(*19*).

Here, we report the first cryo–electron microscopy structure of human TRPM1, which uncovers an unexpected clockwise domain-swapped pore modules that distinguishes it from previously characterized tetrameric ion channels, and provides a structural basis for its constitutive activity. These findings establish a molecular framework for understanding TRPM1 gating and regulation and lay the groundwork for understanding how disease-associated mutations disrupt channel function in visual and pigmentary disorders.

## RESULTS

### Single-Channel activity and constitutive gating of human wild-type TRPM1

The functional characterization of TRPM1 has been hindered by its predominant retention in the endoplasmic reticulum when expressed in heterologous systems(*20*), limiting direct measurements of channel activity using whole cell assays. Consistent with this, Fura-2 AM–based intracellular calcium assays failed to detect TRPM1 activation in response to pregnenolone sulfate (PS), clotrimazole, voriconazole, or capsaicin, despite robust channel expression in HEK293F cells (Fig. 1A). To directly assess channel function, we purified wild-type TRPM1 and reconstituted it into lipid bilayers for single-channel recording (fig. S1A). Using this approach, TRPM1 exhibited spontaneous channel opening events with open probability (Po) of ∼0.7 and a conductance of ∼12 pS in the absence of agonist, indicating constitutive channel activity, as previously suggested but not directly measured (fig. S1E) (*21*). Addition of PS further increased the basal open probability to ∼0.8 and conductance to ∼15.5 pS, consistent with ligand-dependent potentiation rather than obligate ligand-dependent activation (Fig. 1D-E, fig. S1). Notably, the channel activity was inhibited by 2-APB, distinguishing TRPM1 from the closely related TRPM3 channel, which is insensitive to 2-APB (fig S3A-C). Unlike TRPM3, TRPM1 activation by PS did not require PIP₂, although inclusion of PIP₂ increased Po and stabilized channel openings (Fig. 1D-G), indicating a supportive role for membrane lipids(*22*).

**Fig. 1.**
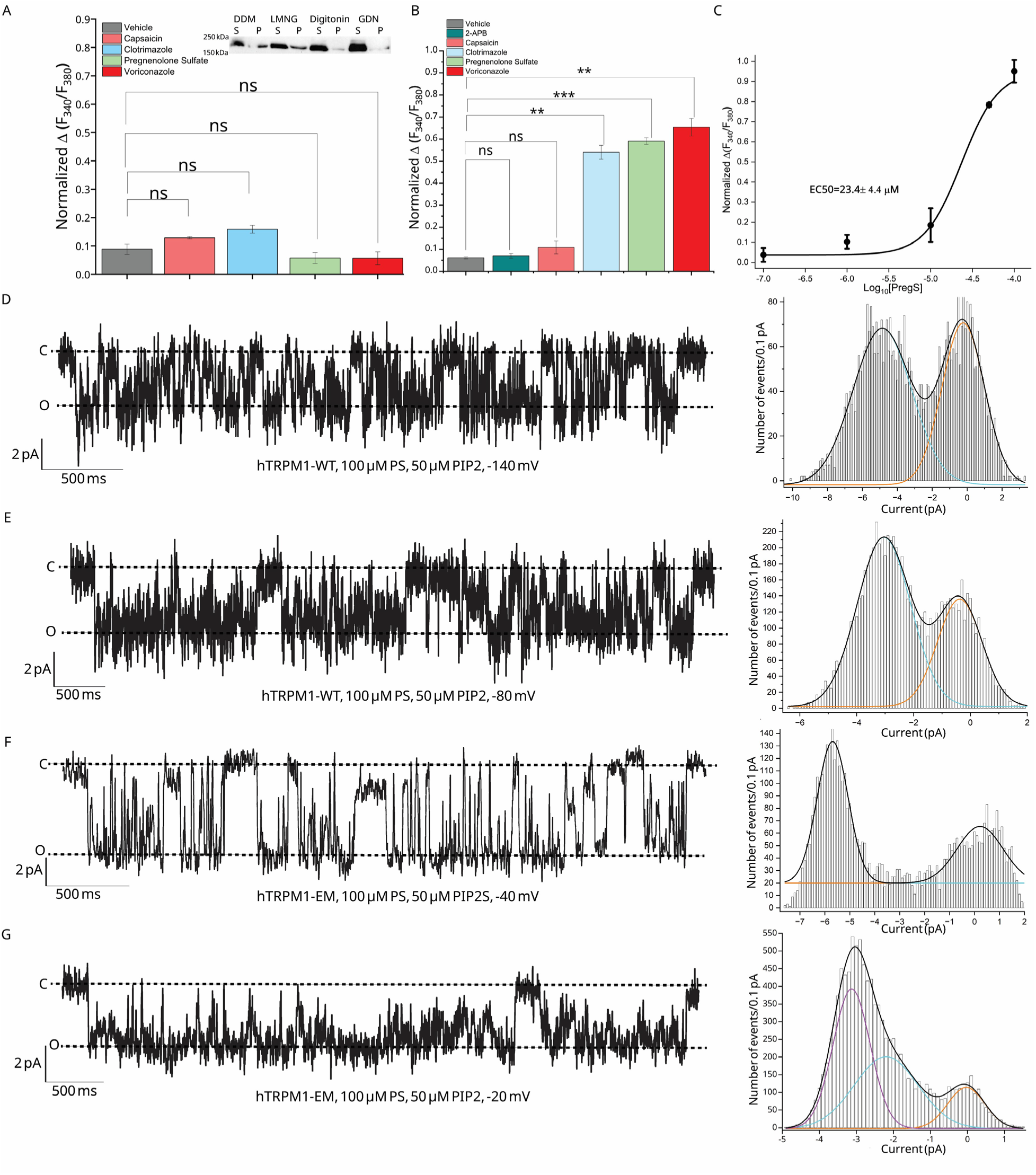
Functional characterization of TRPM1 by Fura-2 calcium measurements and single-channel recordings. Intracellular Ca^²⁺^ responses measured by Fura-2 AM following agonist stimulation of (A) wild-type TRPM1 and (B) the cryo-EM–optimized construct (TRPM1-EM). The inset in (A) shows a Western blot confirming expression of Strep-tagged full-length TRPM1 in HEK293F cells. S and P denote supernatant and pellet (insoluble) fractions obtained after solubilization with the indicated non-ionic detergents. Data are presented as mean ± SEM (n = 3); statistical significance was assessed using a paired two-tailed Student’s t-test. (C) Dose–response analysis demonstrating concentration-dependent activation of TRPM1-EM by pregnenolone sulfate (EC₅₀ = 23.4 ± 4.4 µM). (D-G) Single-channel recordings of purified wild-type TRPM1 and TRPM1-EM reconstituted into planar lipid bilayers are shown in the presence of PregS and PIP2. (D,E) Representative traces and corresponding amplitude histograms for wild-type TRPM1. (F,G) Equivalent recordings and histograms for TRPM1-EM, showing gating behavior between constructs.

Strikingly, single-channel recordings revealed a previously unreported intermediate conductance state (∼1 pA; ∼6 pS at -140 mV) that either persisted or transitioned to a fully open state (∼5 pA; ∼33 pS at -140 mV), consistent with a multistep gating mechanism (Fig. 1, fig S2A). Channel activity exhibited strong voltage dependence, with minimal openings observed at voltages up to −40 mV and a pronounced increase in Po at more negative potentials (Fig. 1D-E, fig. S1). In contrast, channel activity was markedly reduced at positive voltages, revealing a strong gating asymmetry. Together, these data identify TRPM1 as a constitutively active ion channel with discrete conductance states, voltage-dependent gating, and distinct pharmacological sensitivity, defining a gating mechanism that is mechanistically distinct from TRPM3.

## Structure determination of TRPM1

To enable structural studies of TRPM1, we optimized channel expression and stability using fluorescence-detection size exclusion chromatography (FSEC)–based detergent screening for mouse and human TRPM1 orthologs(*23, 24*). Among the two, human TRPM1 exhibited the most favorable expression and monodispersed profiles and was therefore selected for further analysis. To improve sample homogeneity for cryo-EM, we generated a truncated construct lacking exon 11, a flexible poly-lysine loop, and additional disordered residues (hereafter termed TRPM1-EM). This construct displayed markedly improved monodispersity and stability relative to wild-type TRPM1 (fig. S1B).

To determine whether these modifications altered channel function, TRPM1-EM was reconstituted into lipid bilayers and examined using single-channel recordings. Similar to wild-type TRPM1, TRPM1-EM exhibited spontaneous opening events, including a ∼1 pA intermediate conductance preceding transition to a fully open state, and channel activity was potentiated by agonists (Fig. 1F, G, and figs. S1C,G, H & S2D). In contrast to wild-type TRPM1, which required strongly negative membrane potentials for activation, TRPM1-EM opened at less negative voltages in presence of PS and PIP2 (−20 to −40 mV), suggesting that exon11 and/or the poly-lysine loop contribute to the regulation of TRPM1 gating. Because TRPM1-EM localized to the membrane more efficiently than wild-type TRPM1, we evaluated whether it could serve as a robust platform ligand screening and functional characterization of channel activity. Fura-2 AM–based intracellular calcium measurements showed that TRPM1-EM exhibits reliable agonist-evoked Ca^²⁺^ influx, validating its utility for probing TRPM1 pharmacology. In contrast to wild-type TRPM1, TRPM1-EM showed robust Ca^²⁺^ responses upon application of PregS, clotrimazole, and voriconazole, activating the channel with moderate potency (EC₅₀ = 23.4 ± 4.3 µM and 8.3 ± 0.6 µM, 45.3 ±4.4 µM respectively), whereas no activation was observed with 2-APB or capsaicin (Fig. 1B and C; fig. S1C and D). Notably, PregS-evoked Ca^²⁺^ signals were effectively inhibited by 2-APB (EC₅₀ ≈ 40 µM), a compound that does not inhibit the closely related TRPM3 channel (figs. S3 and S4). Together, these results establish TRPM1-EM as a functional surrogate that enables pharmacological interrogation of TRPM1 and reveals ligand sensitivities distinct from related TRPM family members.

To define the overall architecture of TRPM1, we purified TRPM1-EM and subjected it to single-particle cryo-EM. Initial 2D class averages revealed a fourfold symmetric tetramer with prominent intracellular secondary structure features corresponding to the melastatin homology regions, consistent with other members of the TRPM subfamily (fig. S5 and S6). The final three-dimensional reconstruction was resolved to an overall resolution of 4.36 Å (fig. S5B), with higher local resolution observed in the intracellular MHR domains and the C-terminal coiled-coil region (fig. S5 and S6). Several regions, including 80 residues from the N terminus, 211 residues from the C terminus, and few intracellular loops (residues 605-635, 809-853, 1156-1180), were not resolved in the final model, likely due to conformational flexibility.

## Overall structure of TRPM1

The overall cryo-EM density map of TRPM1-EM reveals a broadly conserved architecture within the TRPM family(*15*), with intracellular domains closely resembling those of TRPM3(*16*). In contrast, the transmembrane region exhibits a divergent, noncanonical organization relative to other six–transmembrane (6-TM) tetrameric ion channels.

From the N-terminus, the protein contains the conserved melastatin homology regions MHR1/2 and MHR3/4 (Fig. 2). The MHR1/2 domains form a clamshell-shaped Rossmann-like fold composed of repeating β–α–β motifs, while the MHR3/4 domains are primarily helical, consisting of helix-turn-helix elements that mediate extensive intersubunit contacts. As in TRPM3, these intracellular domains assemble into a large cytosolic platform that supports the transmembrane region. This intracellular architecture is essential for tetrameric assembly and mediates protein– protein interactions that allosterically regulate channel gating including interaction with G-proteins G-proteins (*16, 25, 26*) . Following the MHR domains, as in TRPM3 and other TRPM channels, each TRPM1 subunit contributes six transmembrane helices (S1–S6), with helices S1–S4 forming a voltage sensor–like domain (VSLD) and helices S5–P–S6 assembling into the central ion-conducting pore. The intracellular C terminus begins with the conserved TRP helix, followed by the post-TRP elbow, rib helix, and a central coiled-coil domain that mediates tetramerization (Fig. 2).

**Fig. 2.**
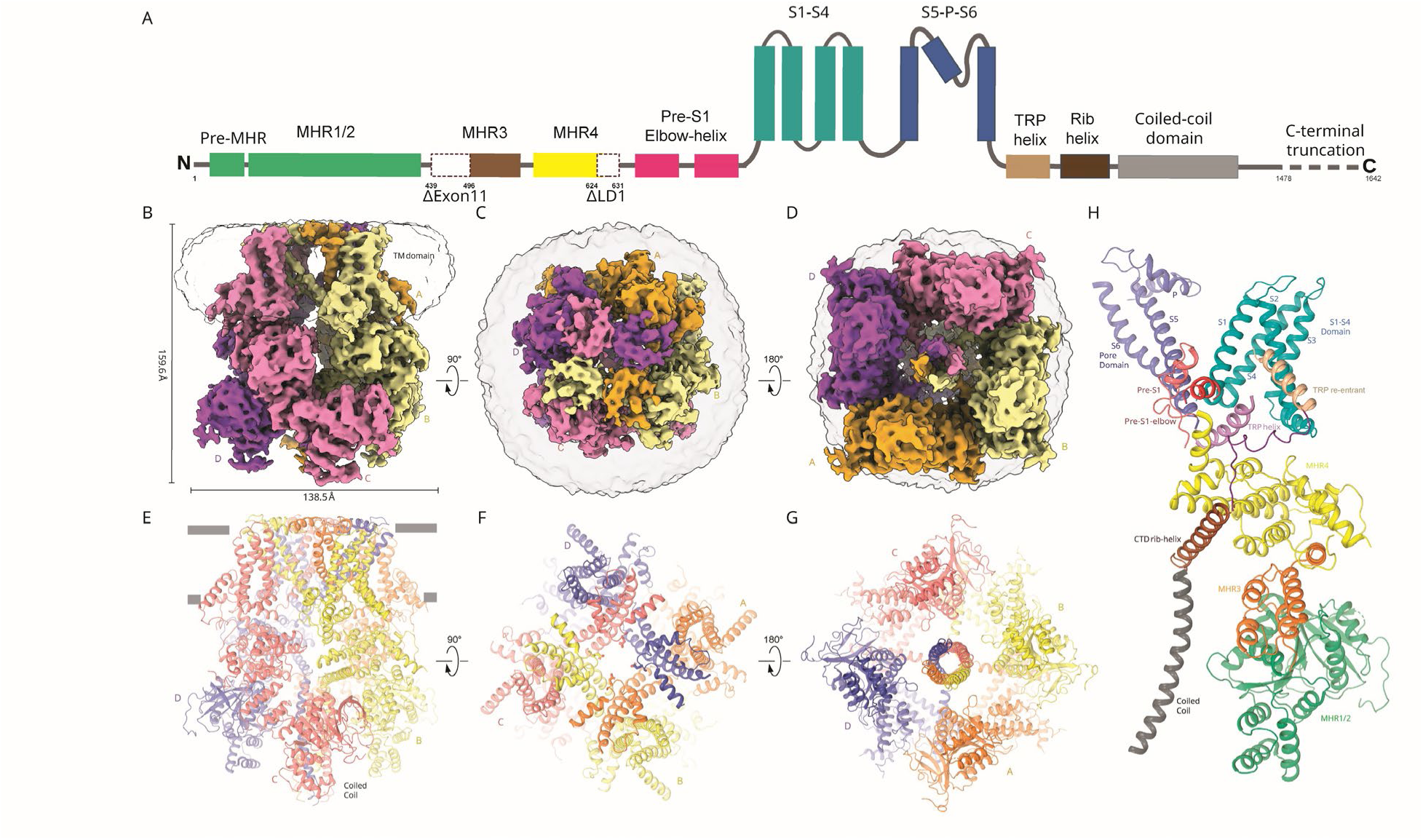
Cryo-EM tetrameric structure of human TRPM1. (A) Linear domain topology of the cryo-EM–optimized TRPM1 construct (TRPM1-EM). Cryo-EM density of human TRPM1 showing in (B) side, (C) top, and (D) bottom views. The corresponding atomic models are shown below each map and colored by subunit (E-G). (H) Structural model of a TRPM1 monomer colored to highlight the major domain architecture.

## Structural discovery of a new transmembrane architecture in tetrameric ion channels

Tetrameric ion channels are thought to have evolved from simple two–transmembrane pore modules into the canonical six–transmembrane (6-TM) architecture, in which an S1–S4 voltage-sensor or voltage sensor–like domain couples to an S5–P–S6 pore module. Across structurally characterized 6-TM channels, including voltage-gated sodium, calcium, and potassium channels, this arrangement is highly conserved, with the pore domain adopting a counterclockwise orientation relative to S1–S4, leading to a stereotyped domain-swapped or non-swapped topology (*6–8, 27*). The cryo-EM structure of TRPM1-EM overturns this prevailing paradigm. While its intracellular assembly conforms to the TRPM family and closely resembles TRPM3, the transmembrane region adopts a fundamentally different organization (fig. S7). Each TRPM1 subunit retains the canonical 6-TM fold, yet their tetrameric arrangement is unconventional: the S5–P–S6 pore module is rotated clockwise relative to the S1–S4 domain, producing a topology opposite to that observed in previously characterized 6-TM tetrameric channels (Fig. 3 and fig. S7 A-E). This reversed orientation defines a previously unrecognized transmembrane fold and demonstrates that alternative domain organizations can support ion conduction.

**Fig. 3.**
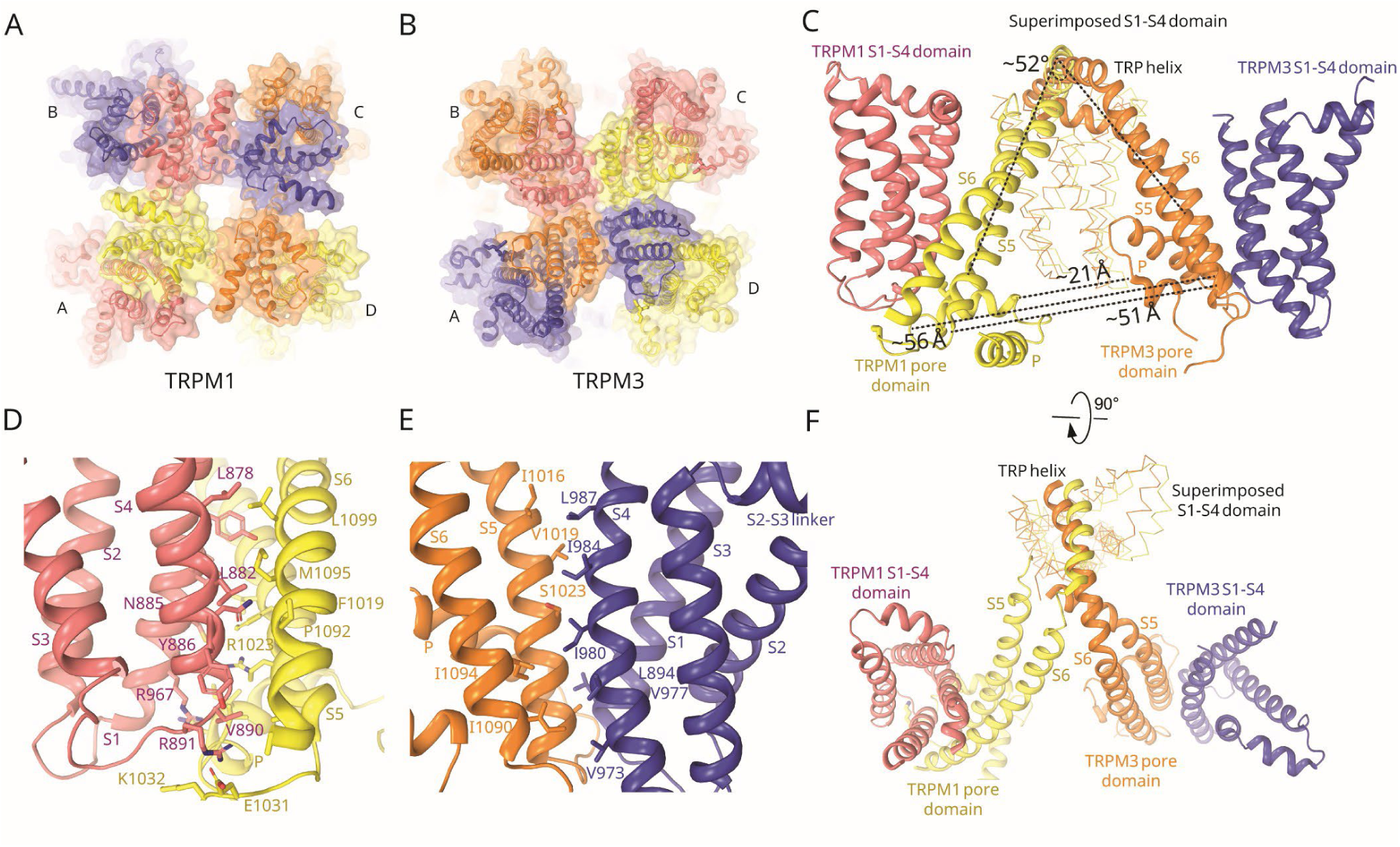
Clockwise pore swapping in TRPM1. Extracellular views of the tetrameric assemblies show pore organization in (A) TRPM1 and (B) TRPM3. (C,F) S1–S4–aligned superposition of TRPM1 (yellow) and TRPM3 (orange), shown after a 90° rotation, reveals opposite pore-module rotations: clockwise in TRPM1 and anticlockwise in TRPM3 relative to the VSLD. Close-up views of pore–VSLD coupling show that (D) TRPM1 forms extensive hydrophobic contacts between the S5/S6 helices and the VSLD paddle, whereas (E) these interactions are different in TRPM3.

Structural superposition of the S1–S4 domains of TRPM1 and TRPM3 reveals that this divergence originates at the S4–S5 junction. Beyond this point, the TRPM1 pore helices undergo substantial translational and rotational rearrangements relative to TRPM3: S5 is displaced by ∼51 Å, S6 by ∼56 Å, and the pore helix by ∼21 Å (Fig. 3). These shifts correspond to an overall clockwise rotation of ∼53° of the pore module, with S5 and S6 rotating ∼50° relative to S1–S4 domain (Fig. 3). Importantly, this reorganization of pore module is not a rigid-body motion. Instead, the pore domain exhibits internal remodeling, including a marked repositioning of S6 relative to S5 that alters helix packing within the conduction pathway. The resulting architecture is stabilized by a distinct network of hydrophobic interactions (Fig. 3D) that differs from TRPM3 (Fig. 3E), highlighting a divergent structural strategy for pore stabilization. Together, these features establish TRPM1 as a structurally unique member of the tetrameric ion channel family, providing a framework for understanding its constitutive activity and distinctive gating behavior while expanding current views of how 6-TM channels can be assembled and regulated.

## Ion conduction pathway in the TRPM1 pore

To define the functional state captured in the TRPM1 structure, we calculated the pore profile and radius at both the selectivity filter and intracellular gate. The selectivity filter is formed by G1056 and the negatively charged E1052 (Fig. 4B), mirroring the conserved architecture observed in TRPM3 (Fig.4A) (*16, 28*). Notably, however, the TRPM1 selectivity filter adopts a markedly wider conformation, with a minimum diameter radius of ∼8.35 Å, compared with ∼3.65 Å in the closed state of TRPM3, showing large lateral fenestration in open state of TRPM1-EM (Fig. 4A-F, fig. S8). A similarly widened selectivity filter was previously observed in PS-bound TRPM3, where ligand binding near the pore helix stabilizes an open selectivity filter conformation (Fig. 4D, fig. S9). (*28, 29*). Like the PS and CIM0216 bound TRPM3, large lateral fenestrations that would be lined by lipid in the native state are seen in the TRPM1 pore (fig. S8). Consistent with this mechanism, we observe additional density adjacent to the TRPM1 pore helix, suggesting lipid-mediated stabilization of the open selectivity filter (fig. S9C and D).

**Fig. 4.**
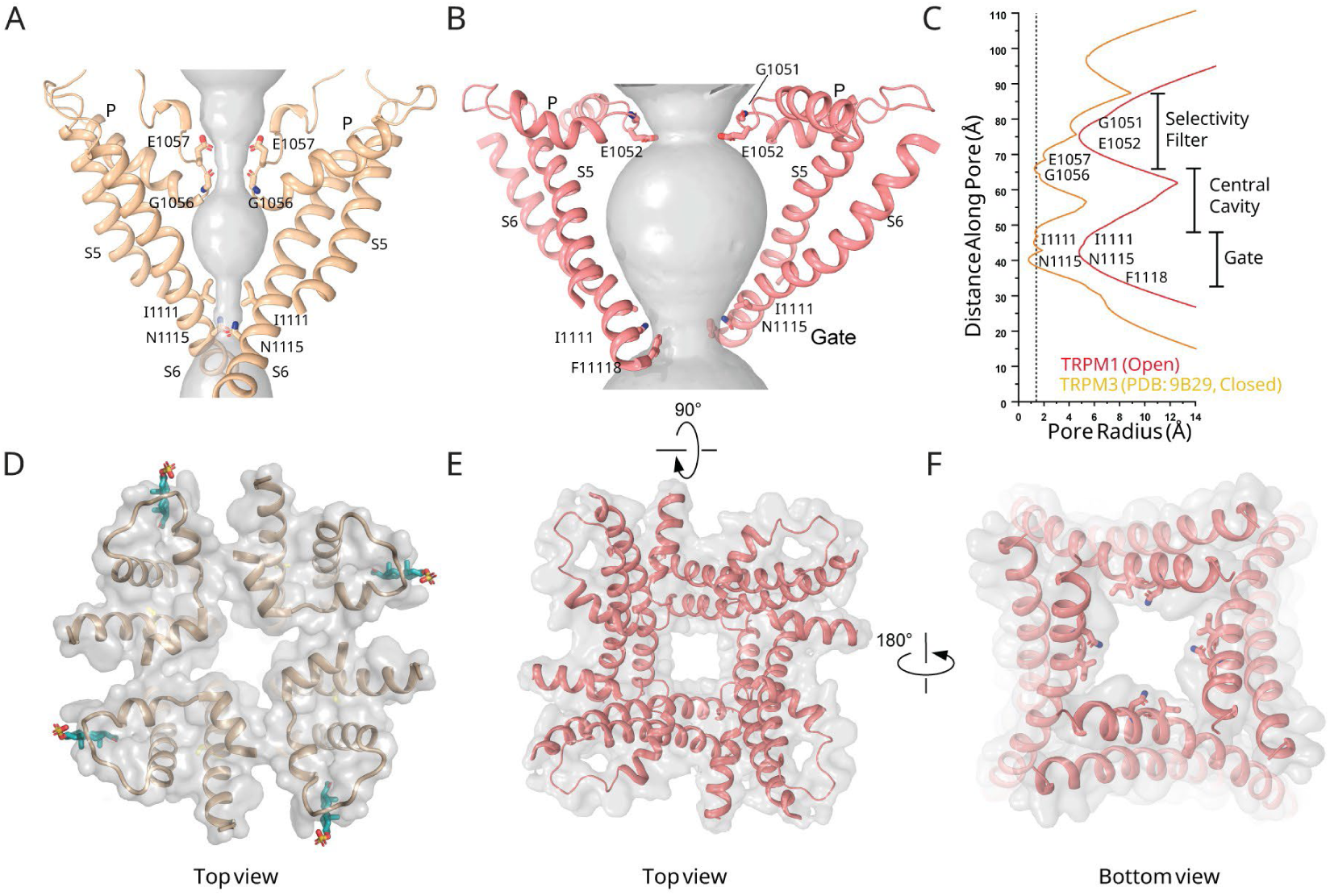
Ion permeation pathway of TRPM1. Surface renderings of the pore illustrate (A) a closed, resting conformation in TRPM3 and (B) an expanded, conducting pore in TRPM1. (C) Pore-radius profiles calculated using the HOLE program compare the constriction points along the ion permeation pathway in TRPM3 and TRPM1. Top views of the selectivity filter show (D) PregS-bound TRPM3 and (E) TRPM1. (F) Bottom view of TRPM1 highlighting the dilated lower gate consistent with an open conformation.

Immediately below the selectivity filter, TRPM1 contains an expanded hydrophobic cavity lined by nonpolar residues, substantially larger than that of TRPM3, potentially accommodating hydrated cations during permeation (Fig. 4B and E). The intracellular activation gate is located at the S6 bundle crossing, where the narrowest constriction is formed by N1116 and I1111. In the TRPM1 structure, this gate is fully open, with a diameter of ∼8.5 Å measured between opposing N1116 side chains (Fig. 4B, C, F). Together, the expanded selectivity filter and fully open gate support the conclusion that TRPM1 is captured in a constitutively conductive state.

## Gating mechanism of TRPM1

The gating architecture of TRPM1 is closely coupled with the unique clockwise rotation of its pore domain relative to the voltage sensor–like domain, which leads to a distinct spatial arrangement of the S5–P–S6 helices. This rearrangement is associated with an expanded selectivity filter compared with TRPM3 and a widened intracellular gate, collectively biasing the channel toward a conductive conformation. The selectivity filter of TRPM1, formed by G1051and E1052, adopts a comparatively open geometry that is consistent with permeation of partially or fully hydrated cations, in contrast to TRPM3, where the narrower filter favors ion dehydration during permeation (Fig. 5A, C, D). In TRPM3 resting closed state, the C-terminal end of the pore helix is positioned deeper within the membrane (approximately 20 Å from the extracellular side) and oriented toward the central hydrophobic cavity, where it contributes to stabilization of permeating ions (Fig. 5A). In contrast, in TRPM1 the C-terminal end of the pore helix lies closer to the extracellular leaflet (approximately 11 Å within the membrane) and adopts a more membrane-parallel orientation rather than projecting toward the cavity (Fig. 5C). This altered geometry is accompanied by an enlarged hydrophobic cavity, likely providing additional space to stabilize hydrated or partially hydrated ions. Notably, a similar rearrangement of the pore helix and selectivity filter expansion has been observed in the pregnenolone sulfate–bound state of TRPM3 (Fig. 5B, fig S9A and B), where channel activation correlates with increased selectivity filter dilation. At the intracellular bundle crossing, the S6 helices diverge substantially, forming a fully open gate lined by N1116 and I1111, with a wide constriction consistent with an activated conformation (Fig. 4B, F and Fig. 5D). S6 exhibits pronounced outward kinking and splays away from the ion conduction axis, further contributing to gate opening (Fig. 5D). Notably, the connection between S6 and the TRP domain adopts a loop-like configuration rather than a continuous helix (Fig. 5H), a feature not observed in other TRPM channels during opening. The S6–TRP helix linker has been shown to adopt distinct conformations across different gating states of TRPM7 and TRPM8, suggesting that this region plays an important role in defining the gating behavior of TRPM family channels.

**Fig. 5.**
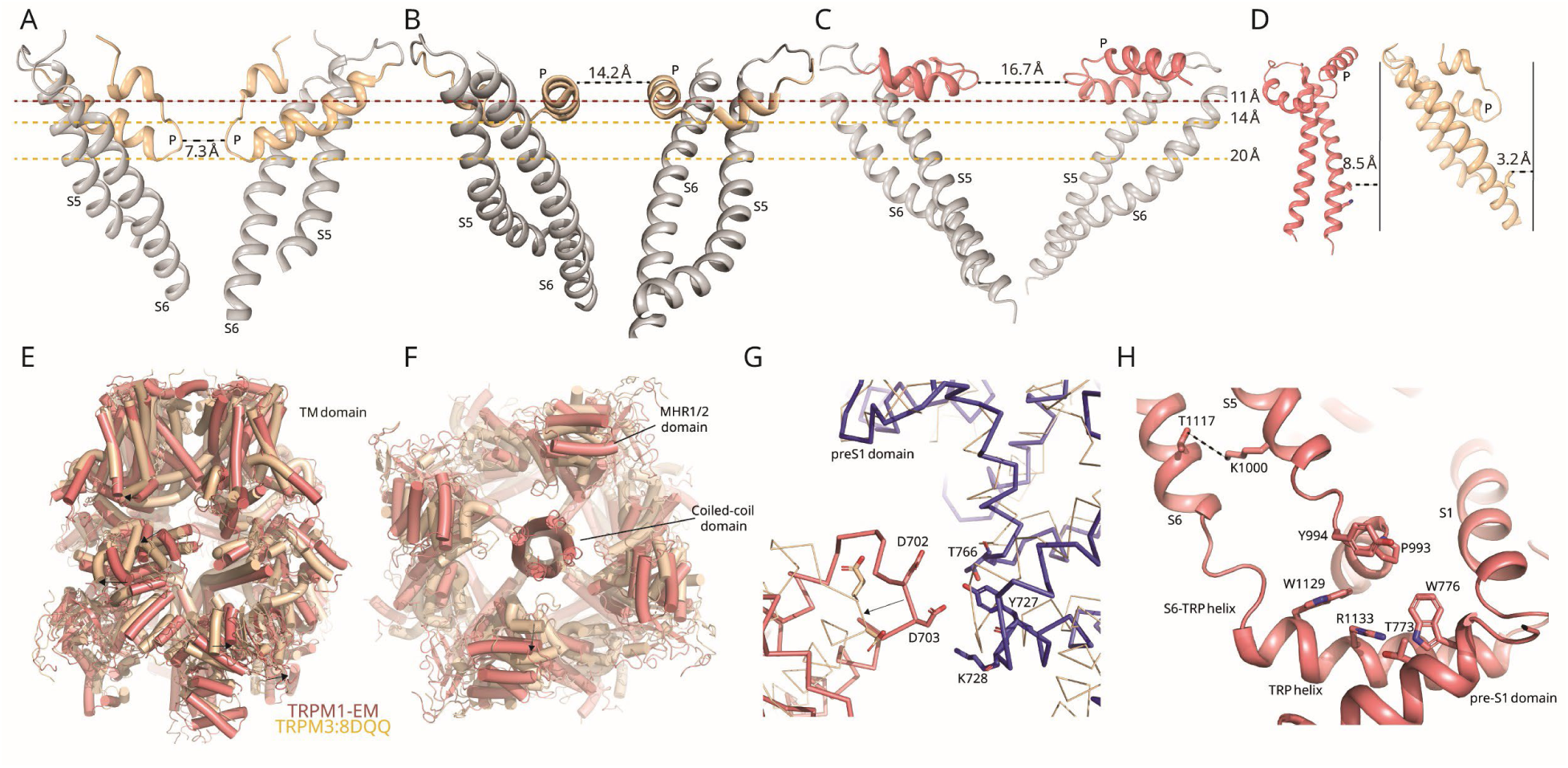
Gating Mechanism of TRPM1. Pore architectures are shown in wheat for (A) the resting closed state of TRPM3 and (B) the PregS-bound state, and in deep salmon cartoon for (C) the open, conducting pore of TRPM1. Lines indicate the relative membrane depth of the pore helix measured from the extracellular side. (D) The S6 segment of TRPM1 forms a continuous α helix positioned ∼8.5 Å away from the ion conduction axis (dark reference line drawn parallel to S6), in contrast to the kinked S6 observed in closed TRPM3. (E,F) Structural alignment of TRPM1 with closed TRPM3, referenced to the S1–S4 paddle domain, reveals coordinated repositioning of the MHR1/2, MHR3/4, rib, and coiled-coil domains, shown in (E) side view and (F) bottom view. (G) The α21–α22 linker within the MHR3/4 domain engages the pre-S1 segment of a neighboring subunit in TRPM1. (H) Structural remodeling at the S6–TRP helix, S4-S5 elbow.

Structural alignment of TRPM1 with TRPM3 based on the S1–S4 scaffold (PDB: 8DDQ) reveals pronounced differences in the relative positioning of the cytoplasmic MHR1/2 and MHR3/4 domains, indicating coupling between intracellular regulatory elements and the transmembrane gate (Fig. 5E and F). More specifically, these comparisons show that, relative to the closed-state TRPM3 structure, the MHR1/2 domain in TRPM1 is rotated upward while the MHR3/4 domain twists toward the transmembrane region. These conformational rearrangements are previously associated with channel activation in other TRPM family members (*29–31*). Consistent with an activated state, the D702 and D703 from α21–α22 motif of MHR3/4 engages the Y727 and T766 of pre-S1 elbow of a neighboring subunit, an interaction characteristic of open-state TRPM4, TRPM7, and TRPM8 structures (Fig. 5G) (*31–33*). These cytoplasmic rearrangements correlate with the expanded ion conduction pathway observed in TRPM1, supporting the conclusion that the structure captures a conductive conformation. Additionally, substantial remodeling occurs at the interface between the S4–S5 elbow, the TRP domain, and the pre-S1 region, where residues W776, Y994, and W1129 adopt altered conformations that mark the pivot point for the clockwise swapping of the S4–S5 linker (Fig.5H). Together, the clockwise rotation of the pore domain, dilation of the selectivity filter, and splayed S6 gate define a gating architecture in which TRPM1 is intrinsically biased toward conduction. This structural configuration provides a mechanistic basis for the constitutive activity of TRPM1 and distinguishes it from other TRPM channels that rely on anticlockwise pore rotation for activation.

## DISCUSSION

In this study, we establish that full-length TRPM1+109 assembles into a bona fide ion channel capable of spontaneous and ligand-modulated activity when reconstituted into planar lipid bilayers (Fig. 1). Our single-channel recordings show that both the native full-length protein and the cryo-EM–optimized construct (hTRPM1-EM) exhibit comparable basal gating and PregS-dependent potentiation, with only modest differences in voltage dependence, indicating that the engineered construct preserves the intrinsic functional properties of the channel (Fig. 1C and D). Although agonist-evoked Ca^²⁺^ influx was detectable only with hTRPM1-EM in Fura-2–based cellular assays (Fig. 1B), this difference is consistent with the limited plasma membrane trafficking and ER retention reported for full-length TRPM1. Thus, the absence of a detectable cellular signal likely reflects expression constraints rather than altered channel function (*20*). Importantly, these findings align with prior work showing that removal of the exon-11 encoded segment improves functional detection of TRPM1 without disrupting channel activity. Notably, TRPM1 was inhibited by 2-APB, in contrast to the closely related TRPM3 channel, which is insensitive to this compound, highlighting a distinct mode of pharmacological modulation (fig. S3 A-C). This differential sensitivity reveals gating features of TRPM1 that were previously difficult to resolve because of challenges in measuring its activity. Together, these findings validate hTRPM1-EM as a faithful structural surrogate and support the conclusion that the cryo-EM structure captures physiologically relevant channel behavior (*34*).

Structural comparison of TRPM1 with other six–transmembrane (6-TM) ion channels reveals a previously unrecognized domain-swapping arrangement in which the pore module is rotated clockwise relative to the voltage sensor–like domains (fig. S7A-F). In contrast, all previously characterized 6-TM tetrameric channels, whether domain-swapped or non-swapped, display a canonical anticlockwise pore orientation (Fig. 6A,B) (*6, 8, 27*). Similar to classical tetrameric channels, the TRPM1 pore forms an inverted teepee-like assembly of S5 and S6 helices that encloses a hydrated central cavity (*6*). Despite its reversed topology, the clockwise arrangement preserves a geometry capable of stabilizing permeant ions within this cavity (Fig.4). The outward orientation of the pore helix and selectivity filter in the clockwise swapped state along with the splaying of S6 segments stabilizes a wide ion conduction pathway in TRPM1 (Fig. 5C), explaining the constitutive activity of the full-length channel observed in our single channel recordings and reported previously (*17, 21, 35*) (Fig. 1).

**Fig. 6.**
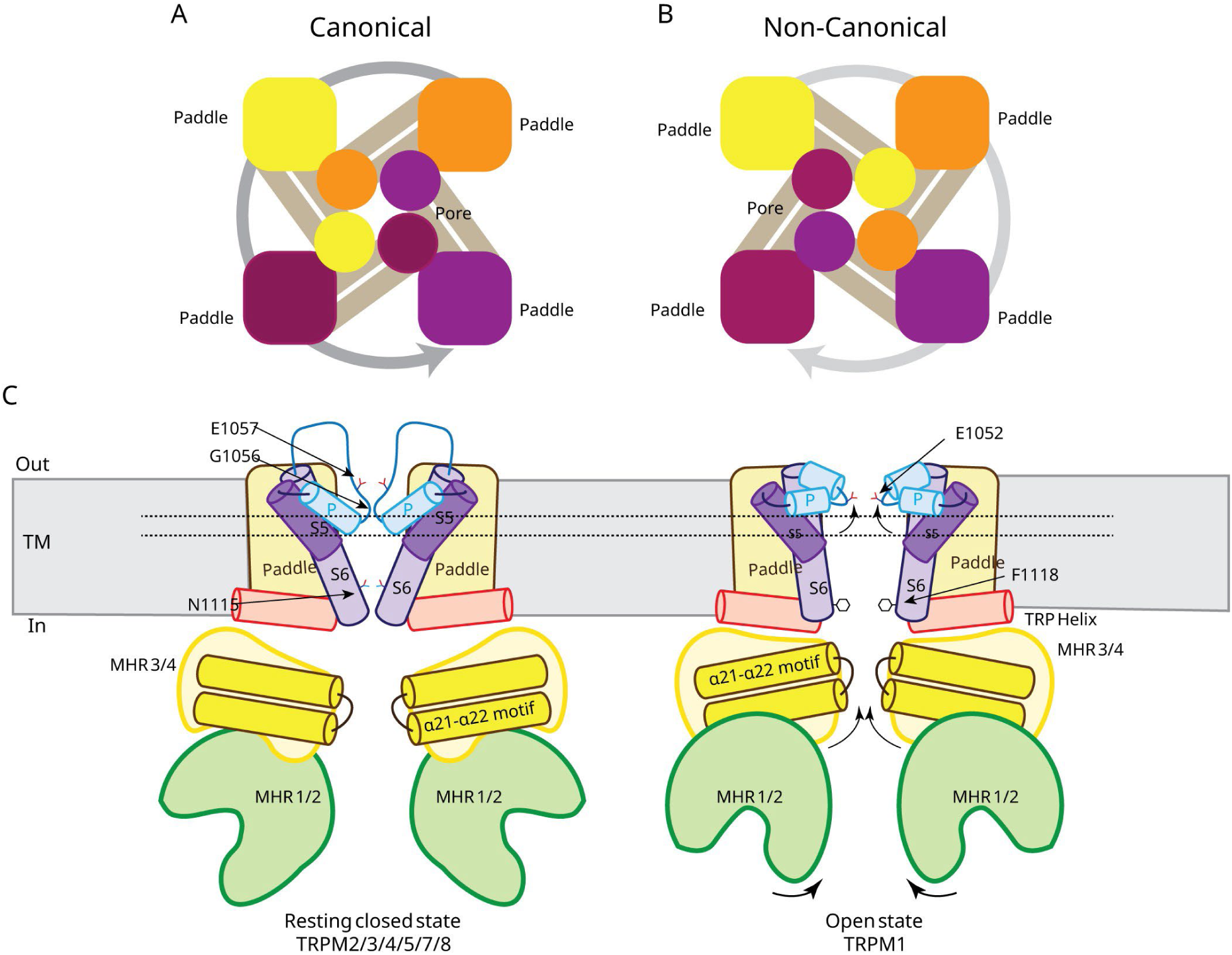
Proposed model for pore architecture and gating in TRPM1. (**A),** Schematic representation of the canonical counter-clockwise pore swapping observed in typical six– transmembrane (6-TM) tetrameric ion channels, where each subunit contributes its pore domain to the adjacent subunit in a counter-clockwise arrangement when viewed from the extracellular side. (**B),** Non-canonical clockwise pore swapping observed in TRPM1. In contrast to the canonical arrangement, the pore domain of each subunit engages the neighboring subunit in a clockwise orientation, resulting in a distinct transmembrane topology and pore architecture. **(C),** Proposed gating mechanism of TRPM1 based on structural comparison with the closed-state structure of TRPM3. TRPM1 adopts a constitutively open conformation stabilized by clockwise pore-domain swapping. Intracellular MHR3/4 elements, particularly residues D702 and D703 within the α21–α22 motif, engage Y727 and T766 located in the pre-S1 elbow of a neighboring subunit. This inter-subunit interaction resembles open-state configurations observed in TRPM4, TRPM7, and TRPM8 (Fig. 5g) and likely contributes to stabilization of the conductive state. Additionally, an upward displacement of the pore helix toward the extracellular membrane side suggests a dynamic pore architecture that may facilitate gating transitions.

Gating in TRPM channels is thought to arise from coordinated intracellular conformational changes that propagate to the lower S6 gate. Structural studies of TRPM4/5/7/8 have identified recurring activation markers, including coupling of the MHR3/4 domain to the voltage sensor–like domain (VSLD) of the cognate subunit and to the pre-S1 elbow of a neighboring subunit, as well as elongation of the lower S6 segment accompanied by shortening of the TRP helix (*30–33, 36*). Although the stimuli that trigger these transitions differ across TRPM family members, the underlying mechanical coupling appears conserved. TRPM1 exhibits several of these activation-associated features despite being purified in an apo state, most notably rotation of the MHR1/2 domain and engagement of the MHR3/4 α21–α22 linker with the neighboring pre-S1 segment (Fig. 6C). These conformational signatures are consistent with stabilization of an open pore architecture and align with the constitutive activity observed electrophysiologically. Importantly, this intersubunit interface provides a structural context for regulation by Gβγ subunits in retinal bipolar cells. Gβγ binds to the exon-17 encoded segment within the MHR3/4 domain conserved between TRPM1 and TRPM3 (*25*), and in TRPM3 such binding disrupts α21–α22 interactions with the pre-S1 region to favor a closed state (Fig. 6B) (*28, 37*). By analogy, Gβγ engagement in TRPM1 is likely to weaken this stabilizing contact, shifting the conformational equilibrium toward pore closure.

Although functional measurements indicate that both TRPM1-WT and hTRPM1-EM are potentiated by pregnenolone sulfate (PregS), the cryo-EM structure determined here under apo conditions captures a pore in an open configuration with a noncanonical architecture. Such an activated conformation in the absence of agonist suggests that TRPM1 intrinsically favors an open-state ensemble, consistent with its constitutive activity observed electrophysiologically. Despite strong sequence conservation and overall structural similarity between TRPM1 and TRPM3 (fig. S4), the clockwise pore-domain arrangement in TRPM1 repositions residues equivalent to those forming the PregS-binding interface in TRPM3, where ligand coordination occurs at the junction of the pore helix (residues W1040 and F1047) and neighboring S1 segment (residues N890 and Y891) in the canonical anticlockwise topology (*28*). In TRPM1, this geometry is disrupted by the reversed domain organization, implying that PregS recognition may occur through an alternative structural configuration. Notably, the pore helix adopts a membrane-parallel orientation reminiscent of activated TRPM3 structures, reinforcing that the captured conformation represents a conductive state (Fig. 5A to D). Determining how PregS engages TRPM1 within this unique architectural framework will be essential for understanding ligand-dependent modulation in channels with noncanonical pore organization and represents an important direction for future investigation.

Finally, disease relevance reinforces the functional importance of this architecture. Mutations in TRPM1 cause congenital stationary night blindness by disrupting signal transmission from photoreceptors to ON-bipolar cells, with missense and nonsense variants impairing channel activity through truncation or destabilization of folding, tetramer assembly, and gating(*38, 39*). Structural mapping localizes multiple disease-associated residues to subunit interfaces and S1– S4–pore contacts, revealing integrity hotspots not apparent from sequence analysis alone (fig. S10). TRPM1 function further depends on its assembly within a bipolar-cell macromolecular complex, where perturbations in associated components likewise compromise channel localization and signaling, suggesting the structural interdependence of this pathway(*40, 41*). Collectively, these observations connect an unconventional pore architecture to constitutive gating and disease susceptibility, establishing a structural framework that links channel organization to retinal pathophysiology.

## Acknowledgments

We thank the ANRF-SERB-Sponsored National Cryo-EM Facility at IIT Kanpur for access to cryo-EM instrumentation. We are grateful to Ravi Ranjan Kumar and Pradeep Kumar Gautam for help with data collection and sample preparation. We also thank Dr. Shikha Singh for valuable comments on cryo-EM processing and for critical suggestions on manuscript preparation. mTRPM3α2 in pCAGGS-IRES-GFP vector was a kind gift from Dr. Stephan Philipp (NCBI Accession No. AJ544535) (*42*).

## Author contributions

MS, KV, and AKS conceptualized the study. Methodology was developed by MS, KV, VK, AC, SN, NV, and RC, while investigation was performed by MS, KV, KM, and AKS. Validation was carried out by MS, KV, and AKS. Visualization was done by MS, KV, and AKS. Funding was acquired by KM and AKS, and resources, project administration, and supervision were provided by AKS. The original draft was written by MS, KV, and AKS, and review and editing were performed by MS, KV, and AKS.

## Competing interests

The authors declare that they have no competing interests.

**Data, code, and materials availability:** The cryoEM density maps and Coordinates have been deposited in the Electron Microscopy Data bank (EMDB) and Protein Data Bank (PDB), respectively, with accession codes XXXX, XXXX.

## MATERIALS AND METHODS

### Construct Design

The human 109+TRPM1 splice variant (accession NP_001238949), originally cloned in pDONR221, was obtained from the DNASU Plasmid Repository (Clone ID HsCD00821818) and subcloned into the pEG-BacMam mammalian expression vector. A structurally optimized TRPM1 construct was engineered to improve biochemical behavior by deleting exon 11 (residues 439–496), removing a polylysine segment (residues 624–631), and truncating the distal C terminus (1479–1642). The pEG-BacMam vector encodes, downstream of the channel C terminus, a thrombin protease cleavage site (LVPRG), followed by an enhanced green fluorescent protein (eGFP) reporter and a Strep affinity tag (WSHPQFEK), enabling fluorescence-detection size-exclusion chromatography screening and affinity purification(*24*). For large-scale expression and structural studies, a non-GFP construct lacking the eGFP fusion was generated. All plasmids were generated using standard molecular cloning techniques and verified by DNA sequencing.

### Protein expression and purification

Recombinant bacmids and baculovirus encoding human TRPM1 (TRPM1), and TRPM1-EM were generated and amplified in Sf9 insect cells (Thermo Fisher Scientific, 11496015) using established protocols (*24*). To produce the baculovirus, Sf9 cells were transfected with approximately ∼21 μg of bacmid DNA using Cellfectin, and baculovirus was harvested after 5–6 days, filtered through a 0.22-μm membrane, and stored at 4°C. For virus amplification, 600 ml of Sf9 cells cultured in SF900 III medium (Gibco, 12659017) were infected and incubated for 88–96 hours, followed by concentration of P2 virus by ultracentrifugation at 63,900g using a T-647.5 rotor (Sorvall WX100, Thermo Fisher). TRPM1 and TRPM1-EM was expressed in suspension-adapted HEK293F cells (Gibco, R79007) cultured in FreeStyle 293 Expression Medium at 37°C with 5% CO₂. Expression conditions were adapted with minor modifications from protocols established for TRPV6(*43*) and GPR158(*44*). Cells were infected at a density of 2.6–3.0 × 10⁶ cells ml⁻¹ with P2 BacMam virus. Sodium butyrate (10 mM) was added 8–10 hours post-transduction, and cultures were incubated for an additional 66–76 hours at 30°C before harvesting by centrifugation at 5,422g for 20 min. Cell pellets were washed once with PBS (pH 8.0) and collected by centrifugation.

All purification steps were performed at 4°C. Cell pellets were resuspended in lysis buffer containing 20 mM Tris-HCl (pH 8.0), 150 mM NaCl, and protease inhibitors (1.5 mM PMSF, 4.5 μM leupeptin, 1.5 μM pepstatin A, and 1.0 μM aprotinin) and lysed by sonication (12 × 10-s pulses, amplitude 20, with 15-s cooling intervals). Cell debris was removed by centrifugation at 9,900g for 18 min, and membranes were pelleted by ultracentrifugation at 186,000g for 1 hour. Membranes were homogenized and solubilized for ∼2.5 hours in buffer containing 20 mM Tris-HCl (pH 8.0), 150 mM NaCl, 1% (w/v) digitonin, and 1 mM β-mercaptoethanol. Insoluble material was removed by ultracentrifugation at 186,000g for 1 hour. The clarified supernatant was incubated with streptavidin resin for 11–14 hours. After washing with 20–30 column volumes of SEC buffer (20 mM Tris-HCl pH 8.0, 150 mM NaCl, 0.01% digitonin), bound protein was eluted with SEC buffer supplemented with 2.5 mM D-desthiobiotin. Eluted TRPM1 and TRPM1-EM was concentrated using a 100-kDa MWCO centrifugal filter and clarified by centrifugation at 40,000g for 30 min. The sample was further purified by size-exclusion chromatography on a Superose 6 10/300 GL column equilibrated in SEC buffer. Peak fractions corresponding to tetrameric TRPM1 and TRPM1-EM were supplemented with 2.5 mM TCEP and concentrated to 0.26 mg ml-1 and ∼2.2 mg ml⁻¹ for single channel recording and for cryo-EM analysis, respectively.

### Cryo-EM sample preparation and data acquisition

Quantifoil R1.2/1.3 holey carbon film grids with a 300-mesh gold support were used for cryo-EM sample preparation as described previously(*45*). Immediately before vitrification, grids were rendered hydrophilic by H₂/O₂ plasma cleaning (10 W, 25 s). Cryo-EM samples were prepared by applying 3 μl of purified TRPM1-EM to the gold side of the grid using an FEI Vitrobot Mark IV maintained at 4 °C and 100% humidity. After a 20-s incubation, grids were blotted for 3 s with a blot force setting of 3 and plunge-frozen in liquid ethane. Cryo-EM data were collected on a Titan Krios G4 transmission electron microscope operated at 300 kV and equipped with a Gatan K3 Summit direct electron detector and a post-column GIF Quantum energy filter in counting mode. Movies were recorded at a nominal magnification of 81,000×, corresponding to a calibrated pixel size of 1.1 Å. A total of 6,752 dose-fractionated movies were acquired across a defocus range of −1.0 to −2.5 μm. Each movie comprised 50 frames with a cumulative exposure of 55 e⁻ Å⁻² collected over 2.0 s, corresponding to an estimated dose rate of ∼33 e⁻ s⁻¹ per physical pixel.

### Image processing and 3D reconstruction

The dataset was processed using RELION(*46*) and CryoSPARC(*47*), or both (refer to Figure. S5, S6, and Extended Data Table 1 for details). A total of 6,752 movies, each comprising 50 frames, were corrected for beam-induced motion using patch motion correction, followed by estimation of the contrast transfer function (CTF) with the patch CTF estimation routine in CryoSPARC (*47*). Following CTF estimation, the exposure curation tool available in CryoSPARC (*47*) was used to reject poor quality micrographs, viz those with a CTF fit value lower than 6 Å, in addition to outliers astigmatism, resulting in 6543 micrographs for further processing. A small subset of micrographs was used initially to select representative particles/templates using blob and template picker, in CryoSPARC, followed by reference-free 2D classification. Finally, a TOPAZ model was first trained against a good set of ∼∼22517 particles encompassing all four classes of TRPM1-EM and then this trained model was used for particle extraction on the entire dataset(*48*). The TOPAZ extraction lead to a total of 3,867,524 particles. Following extraction, the particles were aligned using multiple rounds two-dimensional class averaging and subsequently removing the junk particles; this was followed by ab-initio reconstruction, heterogenous, homogenous, and non-uniform (NU) refinements, resulting in a final subset of 94,538 particles which resolved to a global resolution of 4.36 Å. The reported resolution of the final maps was estimated with the gold-standard Fourier shell correlation (GSFSC) at FSC=0.143 cut-off as implemented in CryoSPARC. The local resolution estimation was done using the local resolution estimation tool in CryoSPARC(*47*). EM density of the reconstructed maps was visualized using UCSF Chimera(*49*).

### Model building and refinement

The cryo-EM reconstruction of the TRPM1-EM achieved an overall resolution of 4.36 Å with the Intracellular domain resolved better compared to transmembrane domain reconstructions. We utilized a model predicted by AlphaFold as a reference to build the intracellular and S1-S4 domain of the hTRPM1 using real-space refinement in COOT(*50*). S5-P-S6 pore domain was built manually with the help of bulky residues as a guide to build the pore domain. The side chain density corresponding to bulky amino acids such as tryptophan, tyrosine, phenylalanine, and arginine helped in correct sequence assignment, with each residue manually adjusted and iteratively built into the cryo-EM density using COOT(*50*). All models underwent real-space refinement against their respective cryo-EM maps with restrained group ADP refinement and were subsequently validated using comprehensive cryo-EM validation in PHENIX(*51*). The UCSF Chimera(*49*), ChimeraX(*52, 53*), and PyMOL(*54*) were used for structure visualization and figure preparation. The cryo-EM data collection parameters and refinement statistics are shown in Extended Data Table 1. Pore radius profile were calculated using HOLE program (*55*).

### Planar lipid-bilayer recordings

Ion channel activity of purified human wild type TRPM1 and TRPM1-EM were examined using planar lipid bilayer recordings as previously described(*56*). A synthetic lipid mixture containing 1-palmitoyl-2-oleoyl-glycero-3-phosphocholine (POPC), 1-palmitoyl-2-oleoyl-glycero-3-phosphoethanolamine (POPE), and 1-palmitoyl-2-oleoyl-glycero-3-phosphoglycerol (POPG) at a 3:1:1 molar ratio was prepared at 30 mM in n-decane. Lipid bilayers were formed across a ∼150-μm aperture of a Meca chip (Nanion) equipped with integrated Ag/AgCl electrodes. Bilayers were separated from symmetric bathing solutions containing 10 mM HEPES (pH 7.4), 300 mM CaCl₂, and 10 mM dextrose. All reagents were of ultrapure grade (>99%, Sigma-Aldrich). Bilayer formation was monitored by capacitance measurements, typically ranging from 5 to 9 pF. For channel reconstitution, 5 μl of TRPM1 (0.002 μg μl⁻¹) was mixed with 50 or 100 μM phosphatidylinositol 4,5-bisphosphate (PIP₂), with or without pregnenolone sulfate, and incubated with 5 μl lipid mixture at 30°C for 30 min before painting onto the aperture. Unitary currents were recorded using an Orbit Mini amplifier (Nanion) controlled by Element Data Reader 3 software. Signals were acquired and digitized at 1.22 kHz. Only recordings containing a single incorporated channel were included in the analysis; traces with multiple channel insertions were excluded. Single-channel events were analyzed using Clampfit together with the Element Data Analyzer (Nanion). All experiments were performed at 20 °C, and data are reported as mean ± SEM.

### Fura-2 Calcium Flux Assays

Full-length hTRPM1 and hTRPM1-EM constructs lacking GFP were expressed in HEK293F cells as described above and performed Fura-AM assay as described previously(*57*). Sixty hours after viral infection, cells were harvested by centrifugation at 300g for 2 min and resuspended in divalent-free HEPES-buffered saline (118 mM NaCl, 4.8 mM KCl, 5 mM D-glucose, and 10 mM HEPES, pH 7.4). Cells were loaded with 5 μg ml⁻¹ Fura-2 AM (Thermo Fisher Scientific) and incubated for 45 min at 37 °C in the dark with shaking at 180 rpm. After a single wash, cells were incubated for an additional 30 min under identical conditions to allow dye de-esterification. Cells were washed twice and resuspended in HEPES-buffered saline supplemented with 2.5 mM CaCl₂ and 1 mM MgCl₂, then maintained on ice in the dark. Fluorescence measurements were performed at room temperature using a Cary Eclipse spectrofluorometer (Agilent Technologies) with continuous stirring. Excitation wavelengths were alternated between 340 and 380 nm at 1-s intervals, and emission was recorded at 510 nm. Intracellular Ca^²⁺^ changes were quantified as the 340/380 fluorescence ratio. Baseline signals were recorded for 1 min before addition of pregnenolone sulfate. Dose–response relationships and EC₅₀ values were calculated using Origin 9.1 (OriginLab).

**Fig. S1.**
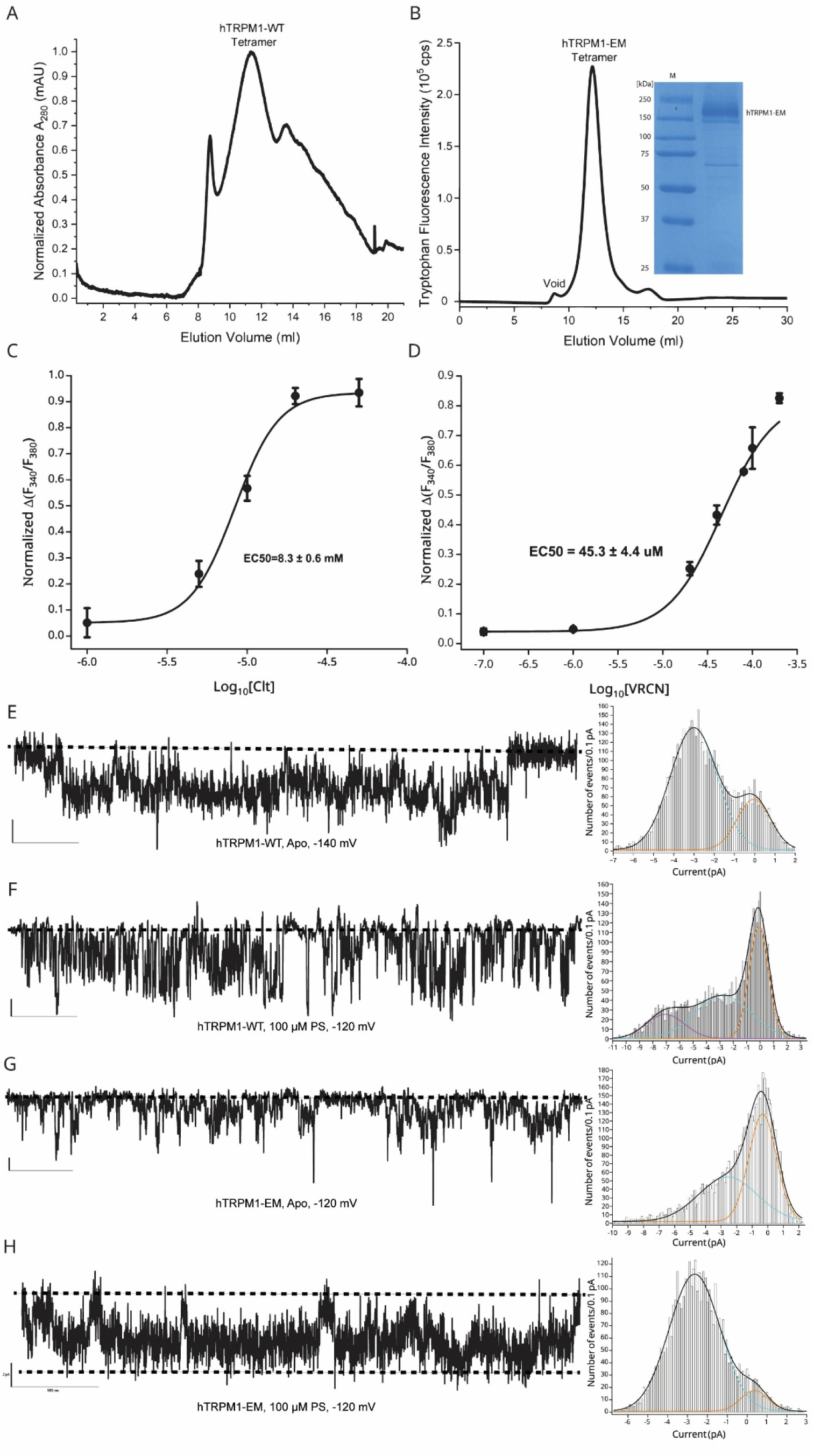
Biochemical and functional characterization of TRPM1 and TRPM1-EM. (A) Size-exclusion chromatography (SEC) profile of full-length wild-type TRPM1. (B) Fluorescence-detection SEC (FSEC) profile and SDS-PAGE analysis of purified TRPM1-EM. (C) Dose– response curves of TRPM1-EM to agonists clotrimazole and voriconazole measured by Fura-2 AM–based calcium flux assays. Each data point represents the average and s.e.m of three independent measurements. Dose response curve through the points were fit with a logistic equation using Origin. (E–H) Representative single-channel recordings and corresponding amplitude histograms for (E) TRPM1-apo, (F) TRPM1 wild-type in the presence of PregS, (G) TRPM1-EM apo, and (H) TRPM1-EM in the presence of PregS. Curve fitting in amplitude histograms were performed in using Gaussian curve fitting module in Origin.

**Fig. S2.**
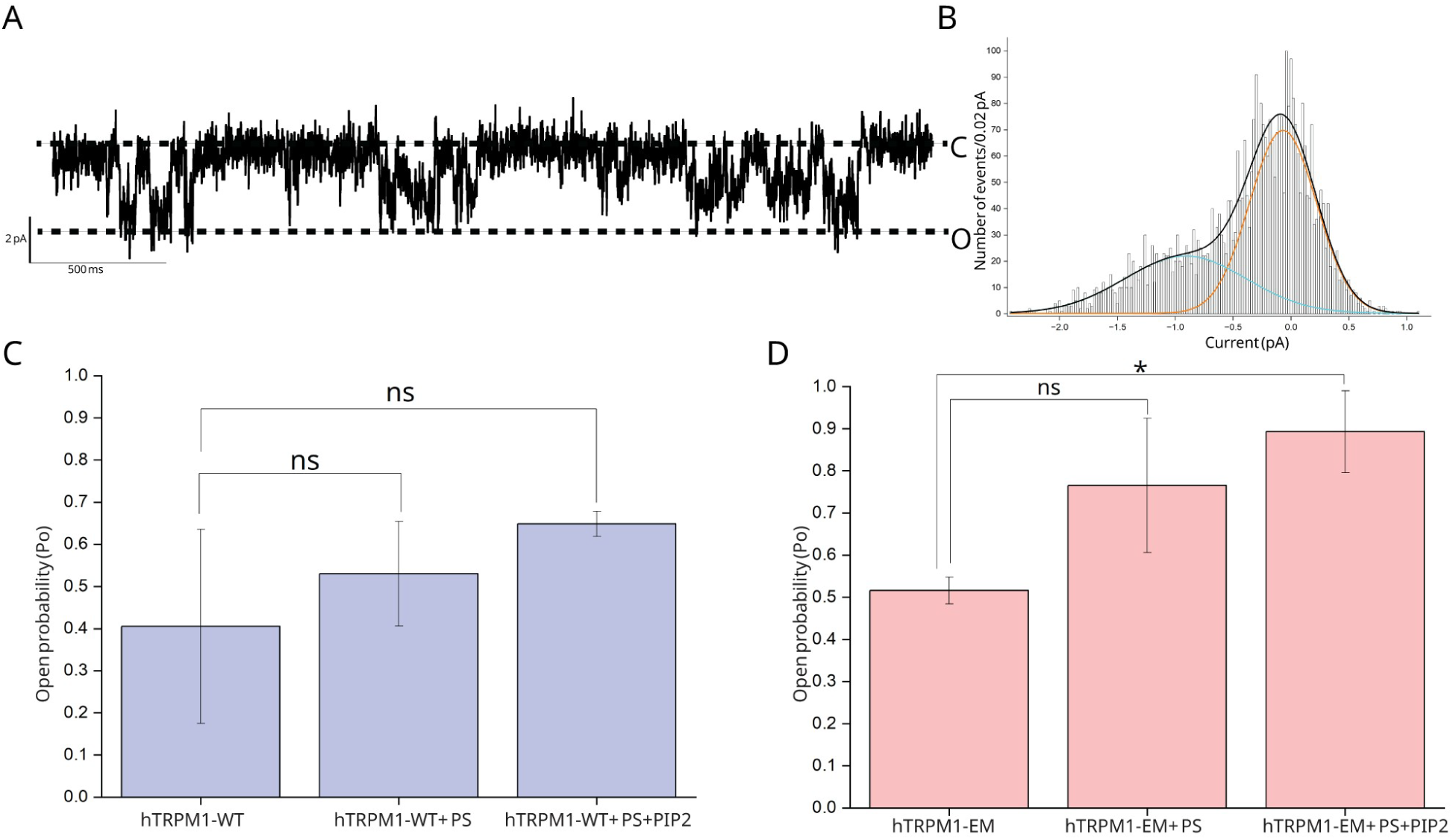
Low conductance states and open probability of TRPM1. (A) Representative single-channel trace showing a low conductance opening of TRPM1-WT in response to PS and PIP₂. (B) Corresponding amplitude histogram of TRPM1-WT channel activity. (C) Quantification of open probability (Pₒ) for TRPM1-WT in apo, PS, and PS + PIP₂ conditions at -140 mV. (D) Open probability of TRPM1-EM under the same conditions. Open probability was calculated as ratio of open channel area (A o) to the total area (A o + A c) in the curve fitted amplitude histograms histograms (*58*). Data are shown as mean ± SEM (n = 3). Statistical significance was assessed using a paired two-tailed Student’s t-test.

**Fig. S3.**
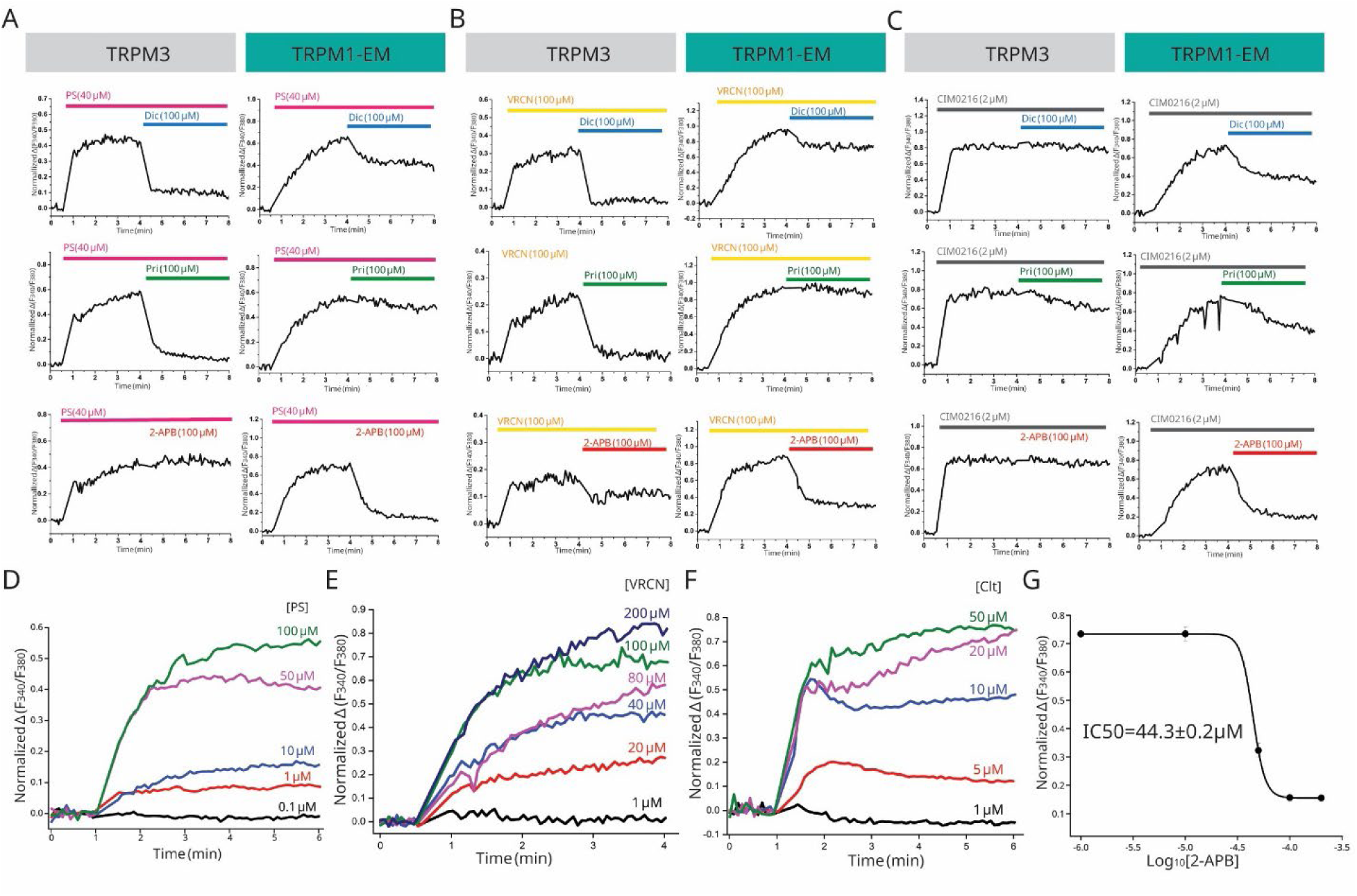
Agonist-evoked calcium responses of TRPM3 and TRPM1-EM and pharmacological modulation. (A–C) Representative Fura-2 calcium imaging traces comparing responses of TRPM3 (left panels, gray header) and TRPM1-EM (right panels, teal header) to different agonists and inhibitors. Cells were stimulated with pregnenolone sulfate (PS; A), voriconazole (VRCN; B), or CIM0216 (C), followed by application of the indicated modulators: diclofenac (Dic), primidone (Pri), or 2-APB, as marked by colored bars. Traces show normalized intracellular Ca^²⁺^ signals (F340/F380) over time. (D–F) Concentration-dependent activation of TRPM1-EM by PS (D), VRCN (E), and clotrimazole (Clt; F), shown as normalized Fura-2 ratio changes. Increasing agonist concentrations produce graded calcium responses consistent with dose-dependent channel activation. (G) Dose–response inhibition curve for 2-APB, plotted as normalized calcium signal versus log inhibitor concentration, yielding an IC₅₀ of 44.3 ± 0.2 μM.

**Fig. S4.**
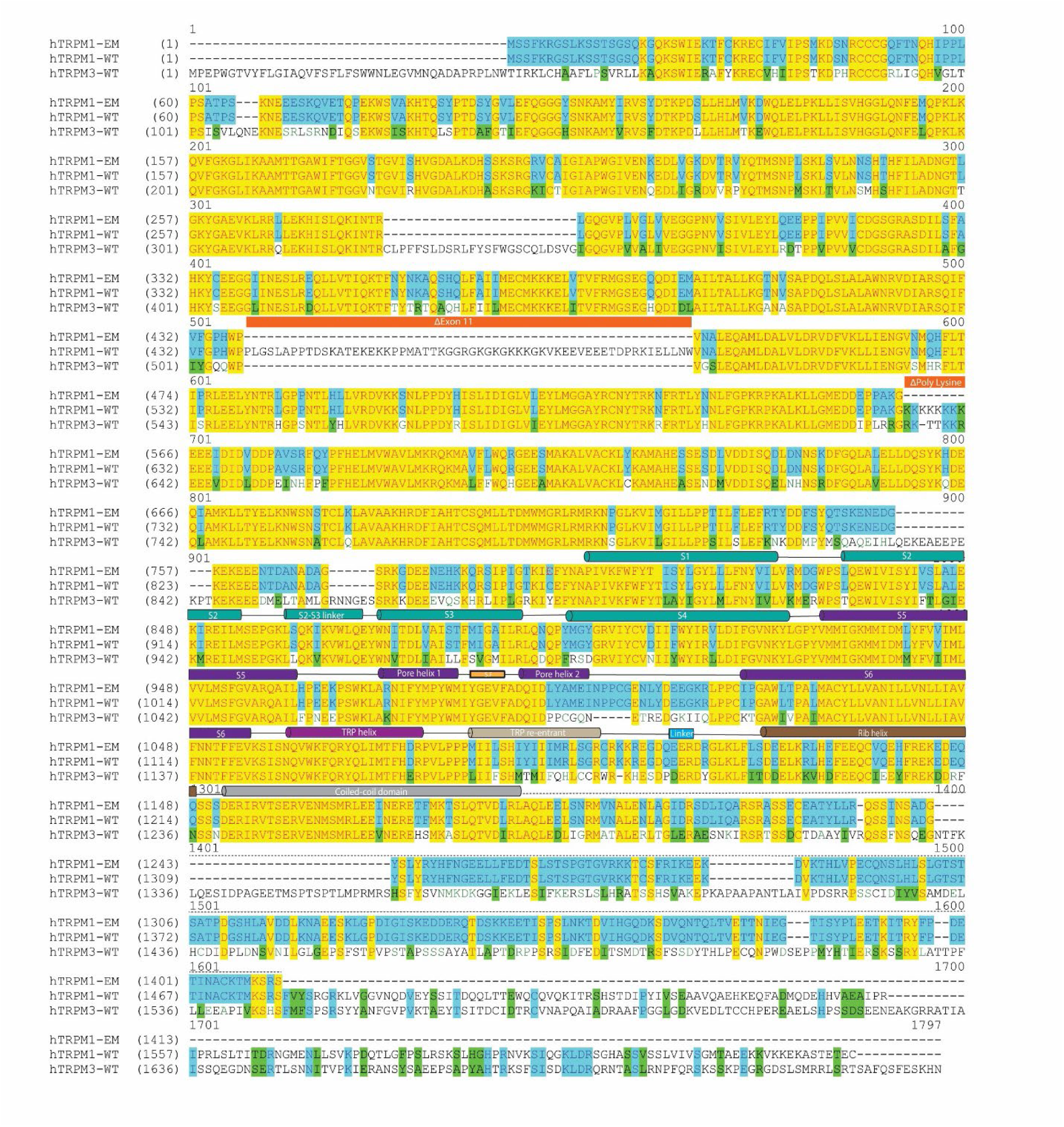
Sequence alignment of human TRPM1, TRPM-EM and TRPM3.

**Fig. S5.**
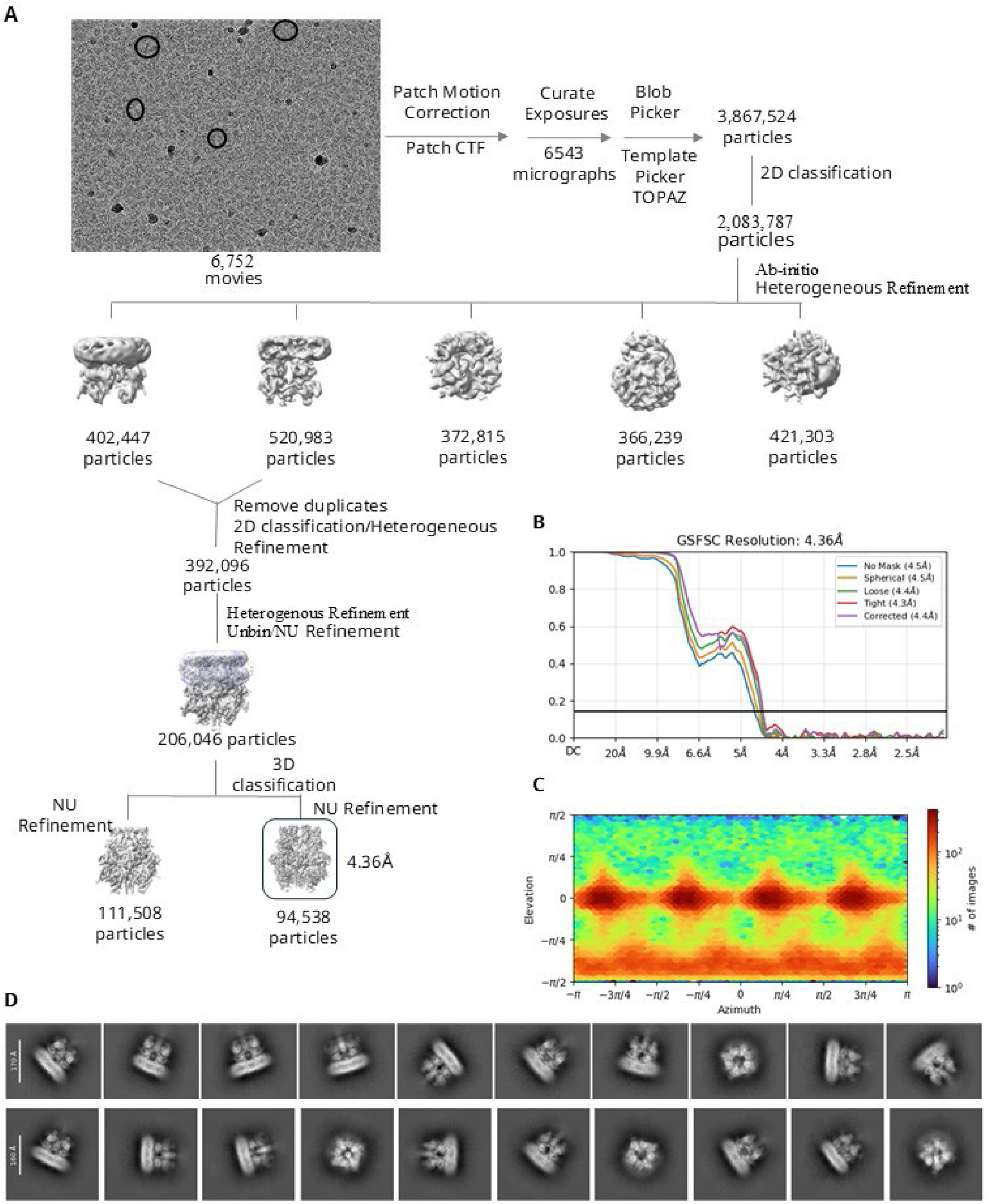
Cryo-EM data-processing workflow for human TRPM1-EM. (A) Representative cryo-EM micrograph of human TRPM1-EM and schematic flowchart of the cryo-EM data-processing pipeline used to obtain the final reconstructions. (B) Gold-standard Fourier shell correlation (FSC) curves for human TRPM1-EM; resolutions are reported at an FSC cut-off of 0.143 (dark gray line). (C) Euler angle distribution of particle orientations for the final reconstruction, calculated in CryoSPARC. (D) Representative 2D class averages of human TRPM1-EM particles.

**Fig. S6.**
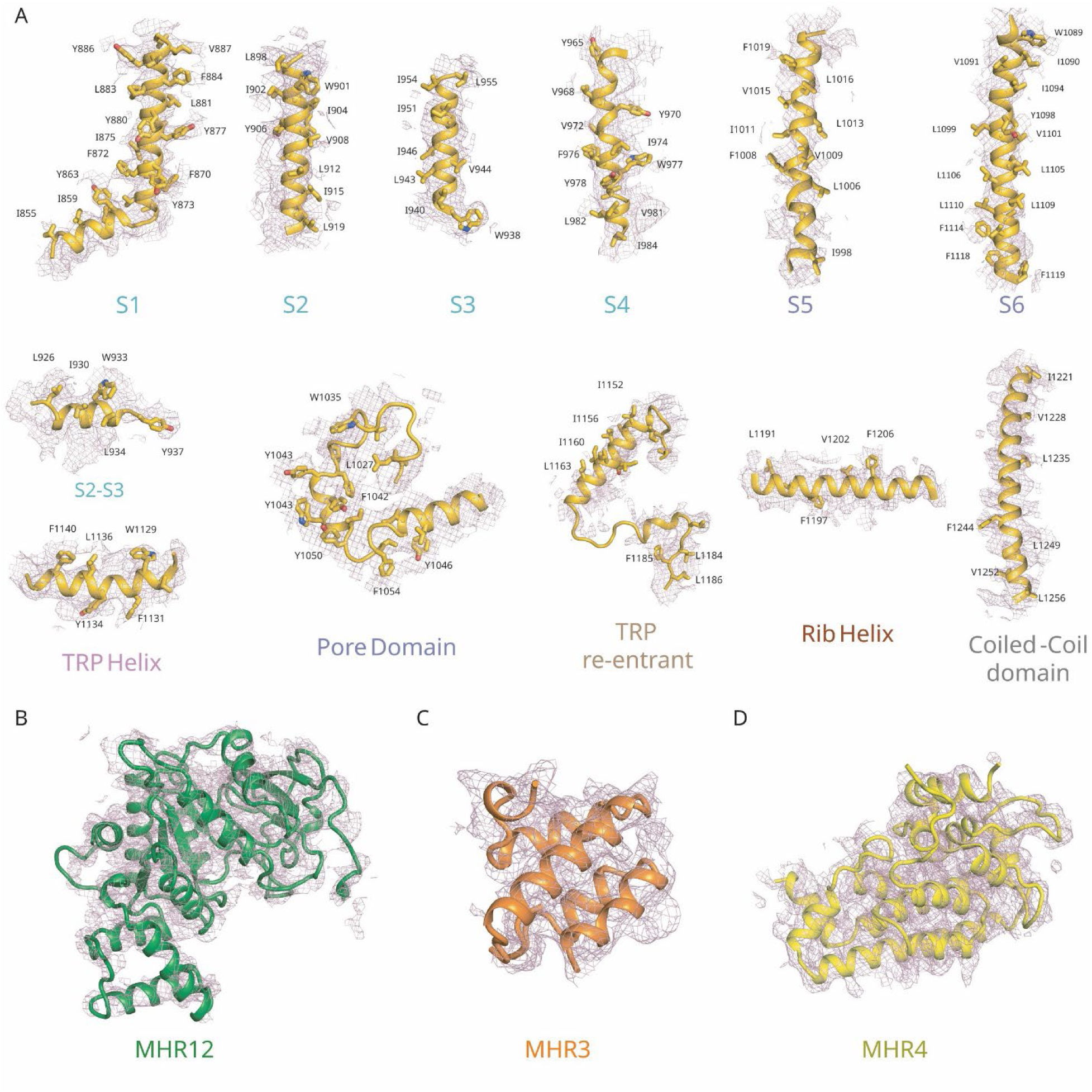
Cryo-EM density and model of TRPM1 secondary structure. (A) Cryo-EM density (mesh) and atomic model of the transmembrane segments and intracellular α-helical regions of each TRPM1 subunit. Cryo-EM density and corresponding models of the (B) MHR1/2, (C) MHR3, and (D) MHR4 domains.

**Fig. S7.**
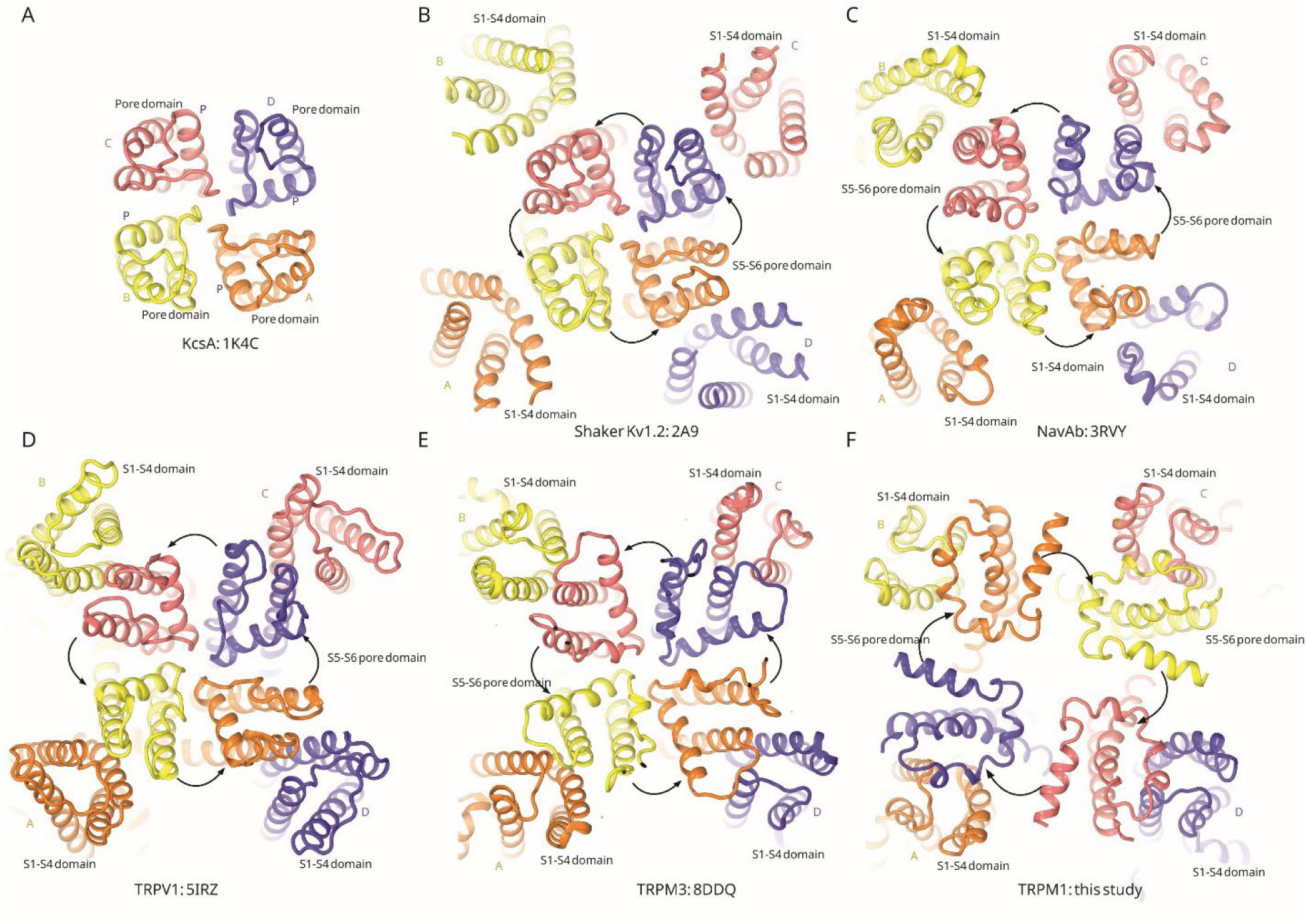
Pore-domain swapping orientations in tetrameric ion channels. Classical two-pore channels, exemplified by KcsA (A), and six–transmembrane (6-TM) cation channels, including Kv1.2 (B), NavAb (C), TRPV1 (D), and TRPM3 (E), exhibit anticlockwise pore-domain swapping when viewed from the extracellular side. In contrast, TRPM1 (F) displays a clockwise pore-domain swap.

**Fig. S8.**
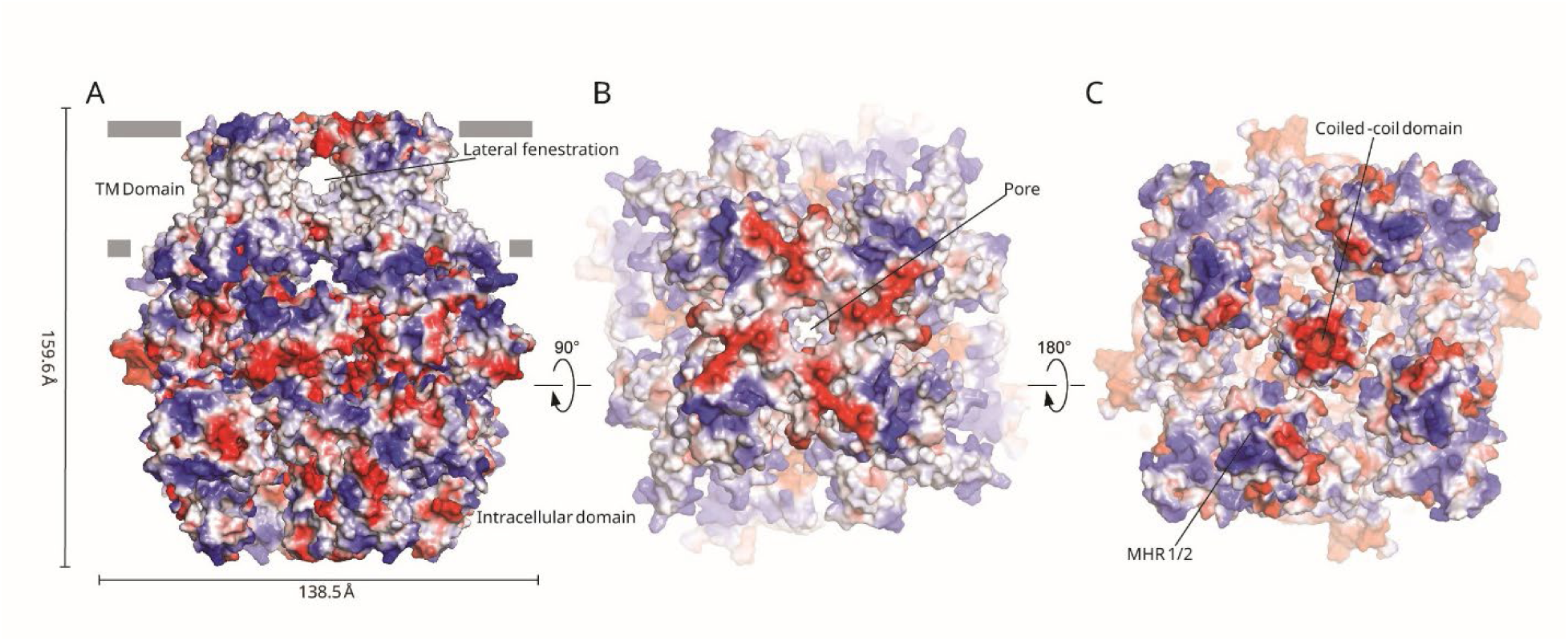
Electrostatic surface potential and lateral fenestrations of TRPM1 in the open state. Electrostatic surface potential of TRPM1 shown in (A) side view, (B) top view, and (C) bottom view. Red and blue indicate negatively and positively charged regions, respectively. Major structural features, including the lateral fenestration, central pore, and coiled-coil domain, are labeled.

**Fig. S9.**
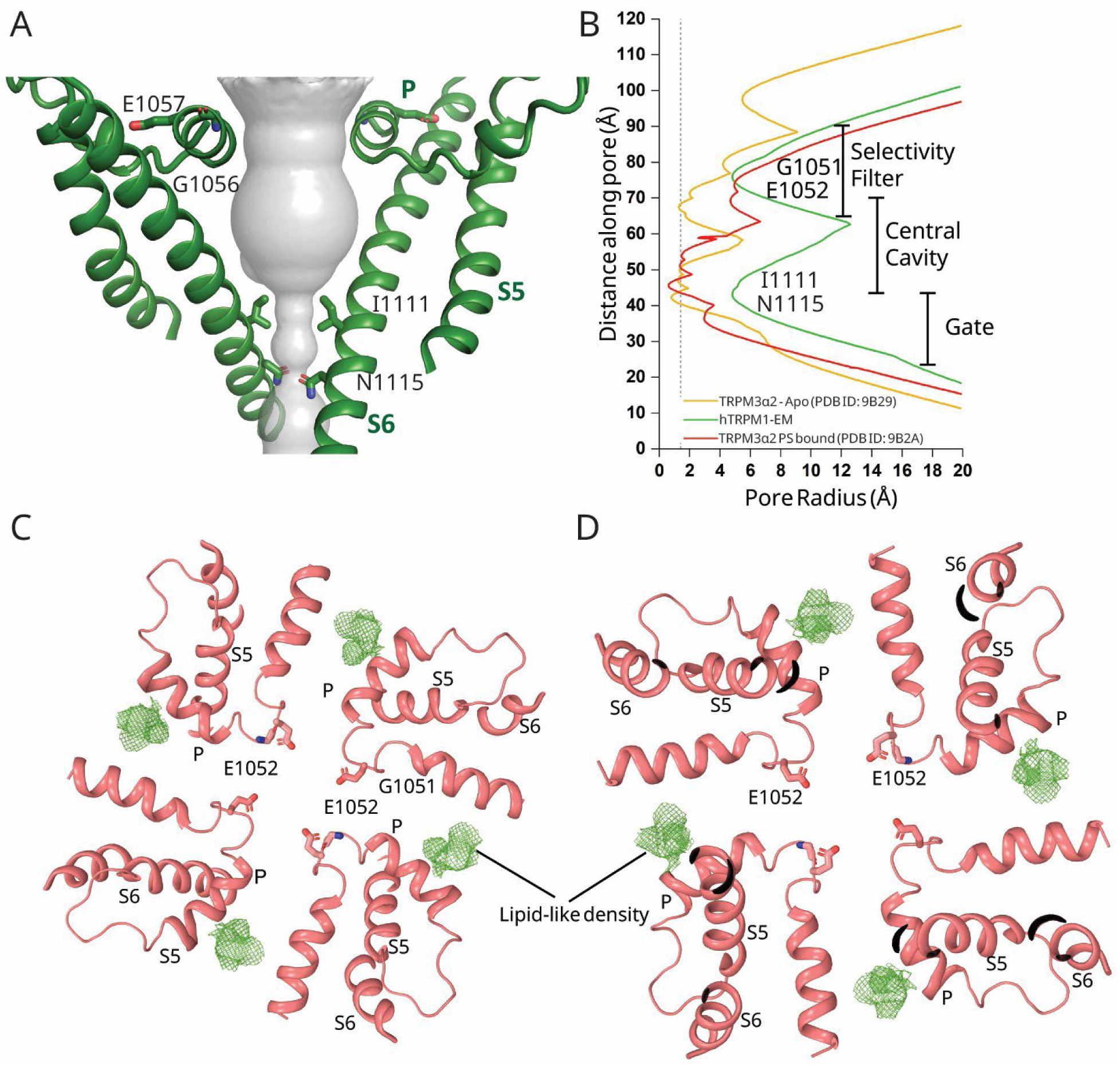
Pore profiles of TRPM1 and TRPM3–PS bound. (A) Surface rendering of TRPM3–PS pore in gray, showing a wider opening at the selectivity filter. (B) Pore-radius profiles calculated using HOLE, comparing TRPM1 with TRPM3 in the apo closed state and in the pregnenolone sulfate (PS)–bound state. (C and D) Unidentified lipid-like density observed near the pore helix of TRPM1, as viewed from (C) extracellular side, (D) intracellular side, reminiscent of the PS-bound TRPM3 structure.

**Fig. S10.**
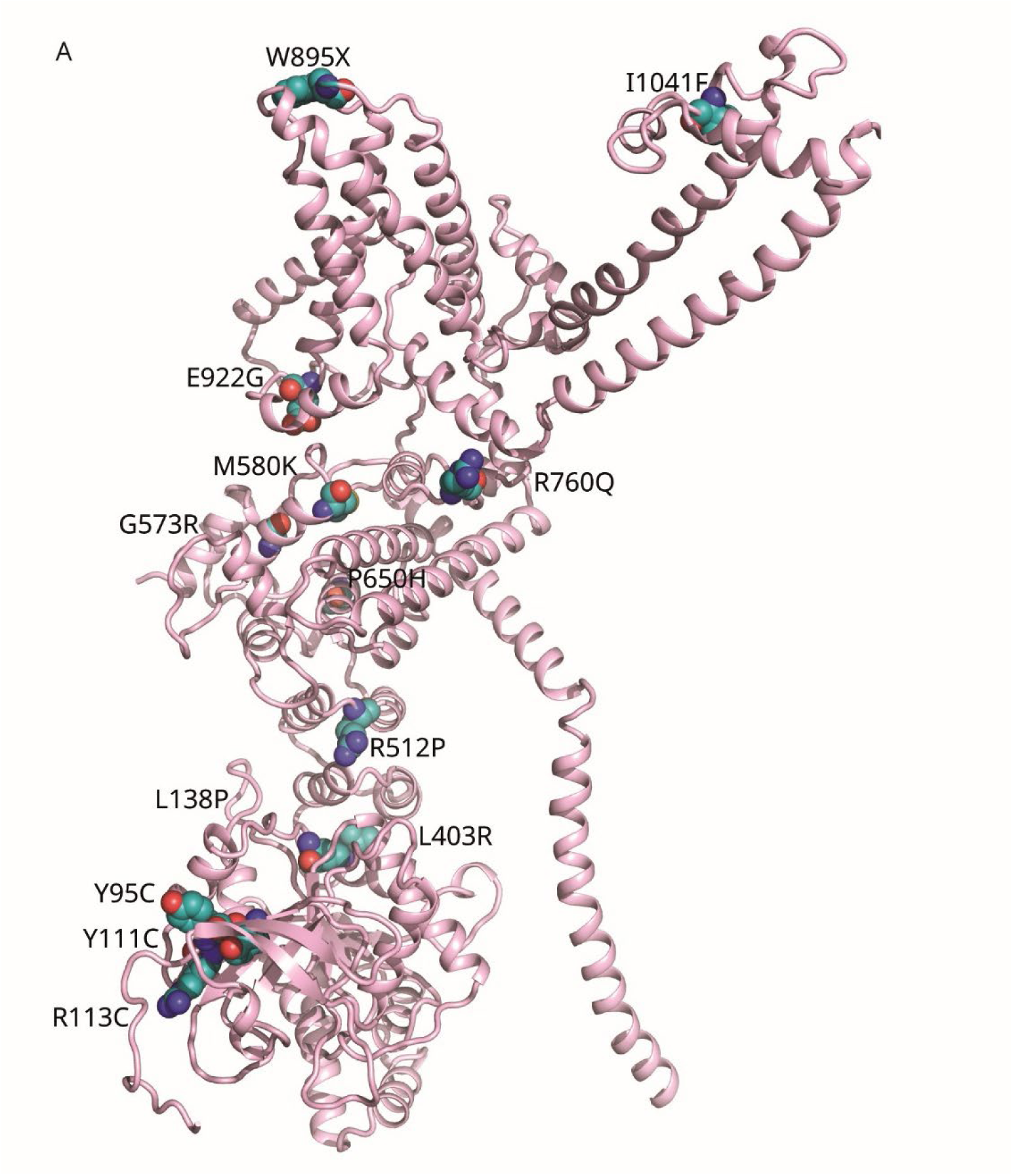
Structural mapping of TRPM1 mutations associated with congenital stationary night blindness (CSNB). Disease-causing TRPM1 mutations, both missense and nonsense, are highlighted as teal spheres and sticks and mapped onto the hTRPM1 structure. This representation provides a structural framework for understanding how specific mutations may disrupt channel folding, gating, or ion conduction.

**Table S1.** Cryo-EM data collection, refinement, and validation statistics.

|  | TRPM1<sub>open</sub><br>(EMDB-xxxx)<br>(PDB xxxx) |
| --- | --- |
| **Data collection and processing** |  |
| Voltage (kV) | 300 |
| Electron exposure (e-/Å<sup>2</sup>) | 55 |
| Defocus range (μm) | -1.5 to -2.5 |
| Pixel size (Å) | 1.1 |
| Symmetry imposed | C4 |
| Initial particle images (no.) | 3,867,524 |
| Final particle images (no.) | 94,538 |
| Map resolution (Å) | 4.36 |
| FSC threshold |  |
| Map resolution range (Å) | 3.1 to 7.0 |
| **Refinement** |  |
| Initial model used (PDB code) | This study |
| Model resolution (Å) | 4.36 |
| FSC threshold |  |
| Model resolution range (Å) | 3.0 to 7.0 |
| Map sharpening *B* factor (Å<sup>2</sup>) | -250 |
| Model composition |  |
| Non-hydrogen atoms | 9,648 |
| Protein residues | 1,131 |
| Ligands | 0 |
| *B* factors (Å<sup>2</sup>) |  |
| Protein | 150.01 |
| Ligand | N.A |
| R.m.s. deviations |  |
| Bond lengths (Å) | 0.011 |
| Bond angles (°) | 1.04 |
| Validation |  |
| MolProbity score | 2.20 |
| Clashscore | 11.8 |
| Poor rotamers (%) | 0.09 |
| Ramachandran plot |  |
| Favored (%) | 88.24 |
| Allowed (%) | 10.91 |
| Disallowed (%) | 0.85 |

**Table S2.** TRPM1 mutations associated with congenital stationary night blindness.

| **S.No.** | **Mutation Type** |  | **Position**<br>**(109 + hTRPM1)** |
| --- | --- | --- | --- |
| 1 | Missense mutation | Loss of Function | Y95C |
| 2 |  | Loss of Function | Y111C |
| 3 |  | Loss of Function | R113C |
| 4 |  | Loss of Function | L138P |
| 5 |  | Loss of Function | L403R |
| 6 |  | Loss of Function | R512P |
| 7 |  | Loss of Function | G573R |
| 8 |  | Loss of Function | M580K |
| 9 |  | Loss of Function | P650H |
| 10 |  | Loss of Function | R760Q |
| 11 |  | Loss of Function | E922G |
| 12 |  | Loss of Function | I1041F |
| 13 | Nonsense mutation | Loss of Function | Q50X |
| 14 |  | Loss of Function | Y813X |
| 15 |  | Loss of Function | W895X |
| 16 |  | Loss of Function | S821X |
| 17 |  | Loss of Function | Y1074X |

